# Fully T2T pedigree assemblies reveal genetic stability and epigenetic plasticity of human centromeres across inheritance and cell-fate transitions

**DOI:** 10.64898/2026.02.14.705860

**Authors:** Shihua Dong, Xiaoyun Xing, Monika Cechova, Hailey Loucks, Selvamani Vijayalingam, Amber Neilson, Monica Sentmanat, Juan Macias, Tianjie Liu, Zheng Dong, Benpeng Miao, Wenjin Zhang, Chad Tomlinson, Heather Schmidt, Edward A. Belter, Ming Hu, Xiaoxia Cui, Nathan O. Stitziel, Karen H. Miga, Ting Wang

**Affiliations:** Department of Genetics, Washington University School of Medicine, St. Louis, MO, USA; Edison Family Center for Genome Sciences and Systems Biology, Washington University School of Medicine, St. Louis, MO, USA; McDonnell Genome Institute, Washington University School of Medicine, St. Louis, MO, USA; UC Santa Cruz Genomics Institute, University of California, Santa Cruz, CA, USA; Genome Engineering & Stem Cell Center (GESC@MGI), Department of Genetics, Washington University School of Medicine, St. Louis, MO, USA; Department of Medicine, Washington University School of Medicine, St. Louis, MO, USA

**Keywords:** Centromere, telomere-to-telomere assembly (T2T), α-satellite, higher-order repeats (HOR), centromeric dip region (CDR), DNA methylation, Fiber-seq, epigenetic inheritance, induced pluripotent stem cells (iPSCs), neural differentiation, *de novo* mutation, X-chromosome inactivation

## Abstract

Centromeres are essential chromosome components yet remain poorly understood due to their highly repetitive sequence architecture. Using fully-phased telomere-to-telomere diploid assemblies from a three-generation pedigree integrated with long-read epigenomes from matched peripheral blood mononuclear cells, induced pluripotent stem cells, and neural progenitor cells, we generate allele-resolved single basepair resolution maps of centromere genetic and epigenetic dynamics across inheritance, reprogramming, and differentiation. We show that centromeric dip regions (CDRs), which define the functional core of centromeres, are positionally stable across generations and cell-fate transitions. In contrast, CDR epigenetic architecture is highly dynamic. Reprogramming markedly attenuates CDR hypomethylation, which is partially restored during differentiation in parallel with global hypomethylation of active alpha-satellite arrays and coordinated changes in nucleosome organization and protein occupancy. Centromeric remodeling is insulated from X-chromosome status, including Xa, Xi, and erosion. Finally, de novo mutations arising during reprogramming are enriched in centromeric regions but depleted within functional centromeric cores.

## Introduction

Accurate chromosome segregation during mitosis and meiosis depends on the centromere, a specialized chromosomal component that nucleates kinetochore assembly and ensures faithful attachment to spindle microtubules^1^. In humans, centromeres are composed primarily of megabase-scale arrays of α-satellite DNA organized into higher-order repeats (HORs)^2^, within which the histone H3 variant CENP-A epigenetically specifies centromere identity^3,4^. Despite their essential role, centromeres have remained among the least understood genomic regions because their extreme repetitiveness has impeded sequence assembly and allele-resolved mapping of sequence and chromatin features.

This landscape changed with the advent of long-read sequencing and telomere-to-telomere (T2T) genome assembly^5^. The complete T2T-CHM13 reference resolved centromeric and other previously inaccessible regions across all autosomes, chromosome X and Y, providing the first comprehensive sequence framework for human centromere biology^6–10^. Building on this foundation, integrative studies have produced complete genomic and epigenetic maps of human centromeres, revealing that α-satellite arrays exhibit structured subdomains and reproducible epigenetic features that correlate with kinetochore organization^10^. In parallel, high-quality genome assemblies from additional individuals from population-scale efforts such as the Human Pangenome Reference Consortium (HPRC)^11,12^ and Human Genome Structural Variation Consortium (HGSVC)^13^ have made it possible to compare centromeres across genomes and begin to quantify the extent of sequence and structural variations that coexist with conserved centromere function. Recent work resolving complete centromeres in multiple human haplotypes further highlights rapid centromere evolution and extensive inter-individual diversity while preserving core functional principles^13^.

Within active HOR arrays, a prominent and recurrent epigenetic hallmark is the localized hypomethylated region, often referred to as centromere dip region (CDR) that coincides with enriched CENP-A chromatin and kinetochore components^7–9^. Importantly, variation in DNA methylation and chromatin organization within CDRs may be associated with changes in CENP-A occupancy and centromere function^14–17^. This observation motivated a model in which hypomethylated CDRs mark functional centromere cores embedded within broader hypermethylated satellite arrays^18^. Recent pedigree-based studies have begun to study the inheritance of centromere stability across generations^19,20^. However, it remains largely unexplored how CDRs themselves behave at the epigenomic level across inheritance, how their epigenetic architecture behaves during major cell-fate transitions, and whether centromeric chromatin is reset and re-established during epigenetic reprogramming. Several fundamental questions remain unanswered in human centromere.

Cellular reprogramming to induced pluripotent stem cells (iPSCs) entails a profound resetting of the epigenome, erasing most lineage-associated DNA methylation patterns and chromatin states^21–23^. While reprogramming-induced epigenetic change has been studied extensively in gene-rich euchromatic regions^24^, centromeric and pericentromeric domains remain much underexplored. This gap is particularly consequential because centromere identity is epigenetically specified by CENP-A, which is known to be misregulated during reprogramming to pluripotency^25,26^. However, whether centromeric DNA methylation architecture exhibits plasticity, or remains insulated from global epigenetic remodeling is unknown. Resolving how these distinct epigenetic layers are coordinated during reprogramming is essential for understanding centromere stability during cell-fate transitions.

Centromeres are embedded within chromosome-scale epigenetic programs. In female somatic cells, X-chromosome inactivation (XCI) establishes a chromosome-wide epigenetic state, and reprogramming is known to trigger heterogeneous erosion of XCI accompanied by widespread DNA methylation remodeling along chromosome arms^27–29^. Importantly, reprogramming is also associated with pronounced X chromosome instability, including frequent aneuploidy, chromosome loss, and ring chromosome formation, highlighting the vulnerability of sex chromosomes during cell-fate transitions^30–32^. However, despite these extensive chromosome-wide epigenetic reconfiguration and instabilities, it remains unclear whether such global remodeling extends into centromeric chromatin, or whether centromeres are insulated from these programs.

Finally, a complementary and clinically relevant knowledge gap concerns genetic stability in centromeric satellites during reprogramming^23,33^. iPSC derivation is known to introduce de novo mutations, raising concerns about genomic integrity in disease modeling and regenerative applications^34–36^; however, mutation burden has largely been assessed in euchromatic regions, leaving the mutational landscape of highly repetitive centromeric satellites largely uncharacterized. Whether different centromeric subdomains—such as active HORs versus inactive α-satellite and pericentromeric repeats—experience distinct mutational pressures remains unknown.

Here we address these questions by integrating fully phased telomere-to-telomere diploid assemblies^20^ with Fiber-seq^37,38^ based long-read, single-molecule epigenomic profiling across a three-generation human pedigree, spanning peripheral blood mononuclear cells (PBMCs), iPSCs, and neural progenitor cells (NPCs). By combining allele-resolved CpG methylation, nucleosome organization and protein occupancy, as well as variant calls with inheritance information, we define centromere architecture, epigenetic remodeling, and mutational burden at unprecedented resolution. Together, our results reveal human centromeres as a stable positional scaffold maintained across inheritance and cell-fate transitions which undergoes centromere-specific epigenetic remodeling during reprogramming and differentiation.

## Results

### 1 Generation and validation of iPSC and NPC lines from a fully phased pedigree to support studies of centromere inheritance

We leveraged a previously characterized three-generation African American pedigree consisting of the grandmother (HG06803; PAN010), grandfather (HG06804; PAN011), their daughter (HG06807; PAN027), and granddaughter (HG06808; PAN028) (**Fig. 1A; Supplementary Table 1**). Fully phased high-quality telomere-to-telomere (T2T) diploid genome assemblies for all four individuals were generated from lymphoblastoid cell lines (LCLs) using multiple long- and short-read sequencing modalities, as described in detail elsewhere^20^, which had been previously validated using Flagger and NucFlag to identify collapsed, duplicated, or otherwise problematic regions, with only ∼0.05–0.15% of bases flagged as potentially erroneous across haplotypes (Supplementary Table 2). Building on this high-quality genomic resource, we reprogrammed PBMCs from all four individuals into induced pluripotent stem cells (iPSCs) using Sendai virus– mediated delivery of the Yamanaka factors (OCT4, SOX2, KLF4, and c-MYC) and subsequently differentiated these into neural progenitor cells (NPCs) to investigate centromere epigenome dynamics during cell-fate transitions (**Fig. 1B; Methods**).

**Fig. 1.**
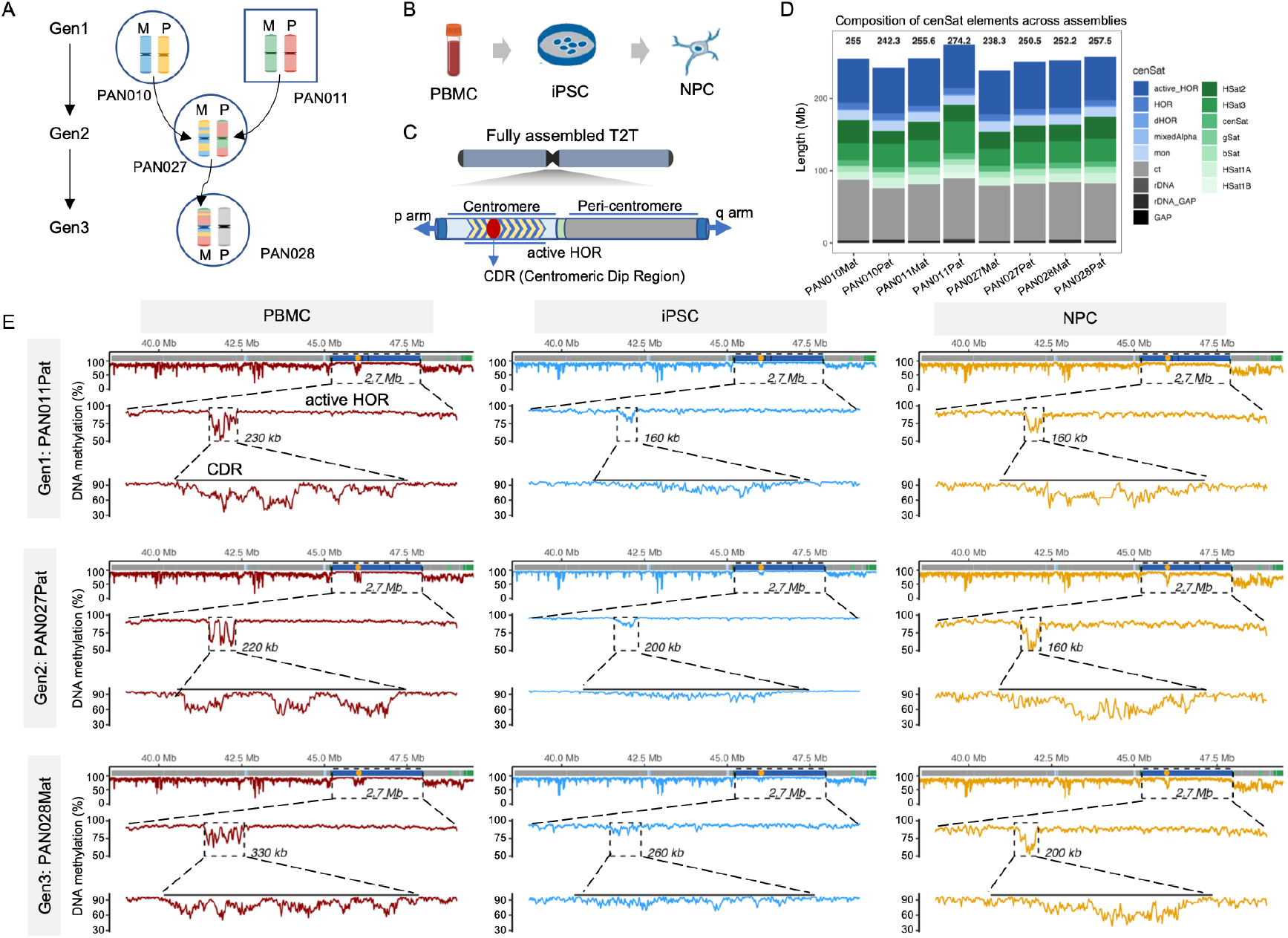
A haplotype-resolved, telomere-to-telomere framework for tracing centromere inheritance and epigenomic remodeling. **A**, Pedigree structure of the three-generation family analyzed in this study. Schematic representations depict the inheritance of chromosome 1 haplotypes across generations. In Generation 1 (Gen1), maternal (Mat) and paternal (Pat) homologous chromosomes are shown in distinct colors, darker shading marks centromeric and telomeric regions. In Generations 2 and 3 (Gen2 and Gen3), inherited haplotypes are shown as mosaics of parental colors, indicating meiotic recombination breakpoints along chromosome 1. Gen, generation. **B**, Experimental design for cell state transitions. Peripheral blood mononuclear cells (PBMCs) derived from individuals of the pedigree were reprogrammed into induced pluripotent stem cells (iPSCs) and subsequently differentiated into neural progenitor cells (NPCs). **C**, Schematic of fully assembled telomere-to-telomere (T2T) genomes used in this study, highlighting the centromere and pericentromeric regions. The active higher-order repeat (HOR) array is indicated, with the centromeric dip region (CDR) defined as a locally hypomethylated domain within the active HOR. **D**, Composition of centromeric satellite (cenSat) elements across haplotype-resolved assemblies. Stacked bars show the total length and relative contributions of different satellite classes, including active HOR, HOR, divergent HOR (dHOR), mixed α-satellite, monomeric α-satellite (mon), centromere transition (ct), ribosomal DNA (rDNA), and gaps, as well as pericentromeric satellites (HSat1A/1B/2/3, β-satellite, and γ-satellite). **E**, Haplotype-resolved CpG methylation tracks across the centromeric satellite (cenSat) domain of chromosome 9 are shown for peripheral blood mononuclear cells (PBMCs), induced pluripotent stem cells (iPSCs), and neural progenitor cells (NPCs) (columns), across generations (rows). For each panel, the top track displays methylation across the full cenSat region. The active higher-order repeat (active HOR) is indicated by a dashed frame, with its length annotated. The colored annotation track above denotes cenSat subdomain composition. Orange dots mark the center position of the centromeric dip region (CDR). Middle panels show a zoomed-in view of the active HOR with the CDR indicated by a dashed frame, and bottom panels further magnify the CDR with flanking 50Kb regions

To enable allele-resolved analysis of centromere features across these cell types, we utilized comprehensive centromere satellite (cenSat) annotation tracks from the reference assemblies^10,20^ in which α-satellite arrays were subdivided into active higher-order repeats (active_HOR), canonical HORs, divergent HORs (dHORs), monomeric α-satellite, and mixed-alpha regions; pericentromeric satellites (HSat1A/1B/2/3, βSat, γSat) and non–α-satellite centromere transition (ct) regions were also annotated (**Fig. 1C, D**). Across chromosomes, peri/centromeric satellite arrays displayed substantial chromosome-specific variation in composition and size (**Supplementary Fig. 1**), but the total centromeric span per haplotype was broadly consistent (Fig. 1E; Supplementary Table 3). This high-resolution centromere map provided the foundation for all subsequent allele-resolved analyses.

To trace centromere inheritance across the pedigree, we identified meiotic recombination points and validated them against independently reported results^20^. Briefly, offspring haplotype assemblies (e.g., PAN027Mat) were treated as references and compared against the corresponding parental haplotype assemblies (e.g., PAN010Mat and PAN010Pat) using dipcall^39^. Genotype similarity between offspring and parental haplotypes was evaluated in sliding genomic windows, enabling identification of large inherited blocks and approximate recombination boundaries. To refine breakpoint localization, homologous sequences flanking each candidate interval were extracted and subjected to local realignment, thereby minimizing interference from genome-wide alignment ambiguity (Methods). Using this strategy, we identified 135 recombination intervals (49 in PAN027Mat, 26 in PAN027Pat, and 60 in PAN028Mat; Supplementary Table 4) (**Supplementary Fig. 2A-C**). Recombination intervals were often sharply resolved, with the smallest interval spanning only 56 bp (PAN028Mat, chr4) (**Supplementary Fig. 2F-G**), and the smallest inherited block measuring 0.96 Mb (PAN027Pat, chr19). Two crossover events were localized within acrocentric chromosomes (PAN027Mat chr13 and chr14) (**Supplementary Fig. 2D-E**), with clearly defined boundaries and one of them precise transmission to the next generation (PAN028Mat) (**Supplementary Fig. 2D**).

Consistent with classical meiotic patterns, recombination events were strongly depleted from centromeric cores. Recombination patterns in PAN027Mat and PAN027Pat were highly concordant with prior reports^20^, while the inclusion of PAN028Mat extends these analyses to a third generation. This independently validated, high-resolution, haplotype-resolved inheritance map enabled base-pair–level tracing of centromere transmission and linking specific centromeric haplotypes to their epigenetic states across cell-fate transitions (**Fig. 1E; Supplementary Table 6**).

Given that cellular reprogramming can introduce genomic rearrangements, we validated the genomic integrity of our derived iPSC and NPC lines using complementary cytogenetic and genomic approaches. First, karyotyping of the original LCL lines confirmed diploid genomes for PAN010, PAN011, and PAN027, whereas PAN028 exhibited a mosaic Turner karyotype (ISCN 45,X[5]/46,XX[15])^20^. Following reprogramming, all iPSC lines were clonal and karyotypically normal (**Supplementary Fig. 3**), including PAN028, in which the mosaicism detected in the original LCLs was resolved; two independent iPSC clones from PAN027 were included in the study.

To enable allele-resolved characterization of centromeric epigenomic features, we generated long-read sequencing data from all iPSC and NPC lines. PacBio Fiber-seq was performed for all samples, yielding 31–73× coverage with a mean read length of 20.6 kb, and Oxford Nanopore sequencing was additionally generated for the two PAN027 iPSC clones (35.2×; Supplementary Table 5). Principal component analysis (PCA) of genome-wide CpG methylation profiles separated samples primarily by cell type rather than by individual, clone, or sequencing technology, with the two PAN027 clones clustering closely together (**Supplementary Fig. 4**), indicating high reproducibility of reprogramming-induced epigenetic states.

All long-read data were independently aligned to the corresponding high-quality, haplotype-resolved assemblies. Reads were phased based on alignment uniqueness, mismatch rates, and alignment scores (Methods). PacBio reads achieved a median phasing rate of 98.75%, whereas ONT reads achieved a lower phasing rate (83.94%), largely due to their shorter and more variable read lengths (**Supplementary Fig. 5A**). Alignment mismatch rates were substantially reduced after phasing, confirming accurate haplotype assignment (**Supplementary Fig. 5B**). Read-depth analysis demonstrated uniform coverage across chromosomes, including centromeric regions, for both maternal and paternal alleles (**Supplementary Figs. 6, 7**). Together, this experimental and analytical framework enabled allele-resolved tracking of CpG methylation, chromatin accessibility, and genetic variation across inheritance, cellular reprogramming, and differentiation (**Fig. 1E**).

### 2 Centromere DNA methylation dynamics during reprogramming and neural differentiation

Centromeric dip regions (CDRs) are localized domains of reduced DNA methylation within α-satellite higher-order repeat (HOR) arrays and have been shown to coincide with regions enriched for CENP-A chromatin. CDRs typically consist of multiple consecutive subCDRs separated by methylated spacers, with each centromere containing a single CDR generally localized within the active HOR array. We identified CDRs as locally depressed methylation valleys within active HORs using the PacBio long read data (Methods) from PBMCs (**Fig 2A; Supplementary Fig. 8; Supplementary Table 7**). To assess consistency with previously established CDR definitions, we separately applied a previously reported CDR-identification approach^20^ to the same PBMC datasets, which defines subCDRs for each chromosome (**Supplementary Table 8**). Across all assemblies and chromosomes, all CDRs identified in this study showed complete overlap with those defined by the previous method, and 98.7% (762/772) of previously reported subCDRs overlapped with our identified CDRs (e.g. Chr 1 in **Supplementary Fig 9**), confirming the robustness and concordance of our CDR identification. Across the pedigree, five independent centromeric haplotypes are present (PAN010 Mat/Pat, PAN011 Mat/Pat, and PAN028 Pat), whereas PAN027 Mat/Pat and PAN028 Mat haplotypes are directly inherited from their parents, providing multiple opportunities to assess CDR stability across both inheritance and cell-fate transitions. Concordant with previous reports in the LCL lines from this pedigree^20^, CDR positions in PBMCs were highly stable across generations (**Fig 2A; Supplementary Fig. 11**).

**Figure 2.**
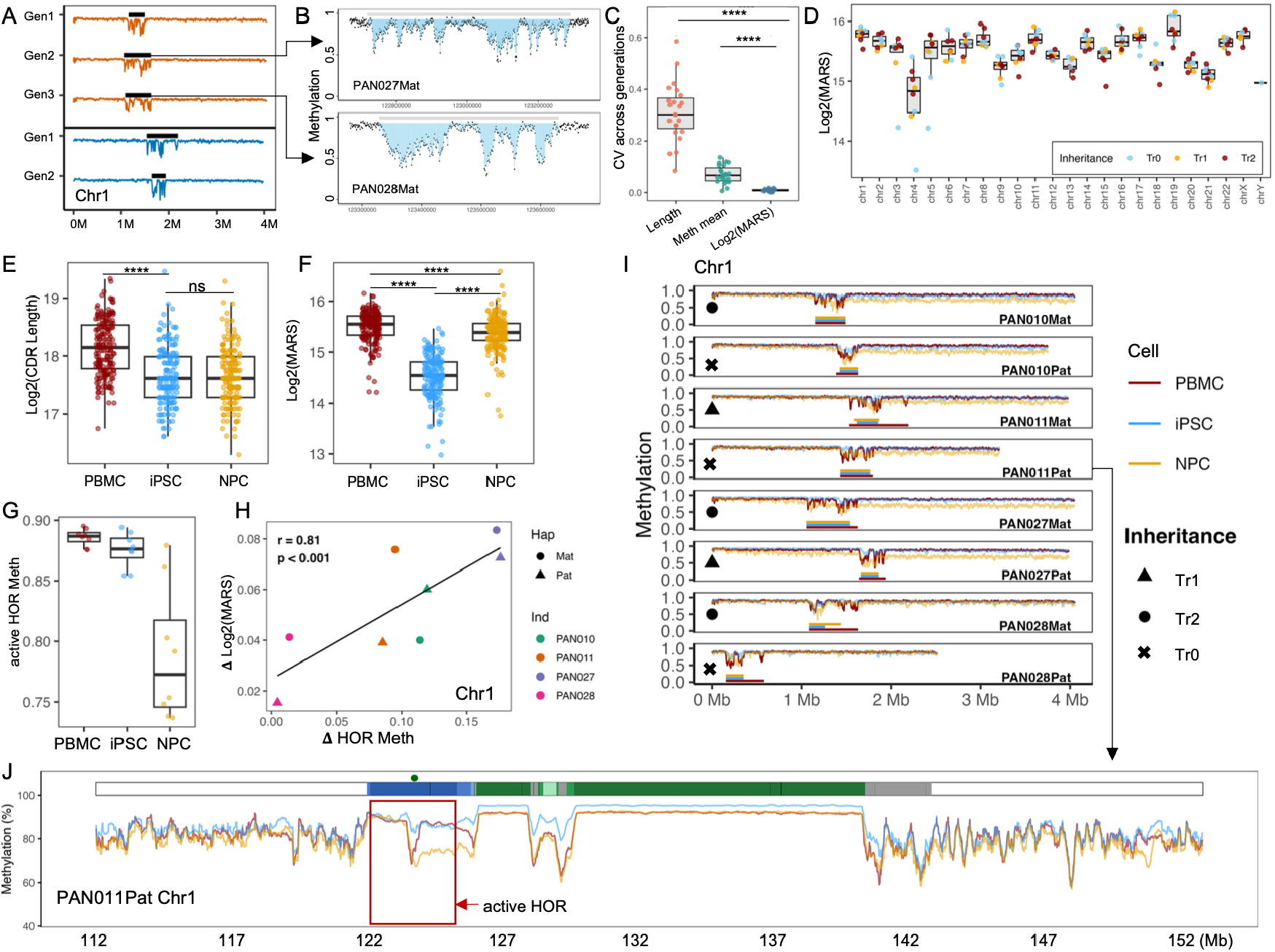
Centromeric dip regions exhibit stable inheritance but dynamic, cell-state–dependent epigenomic remodeling. **A**, DNA methylation profiles across the active higher-order repeat (HOR) array of chromosome 1 illustrating CDR inheritance with coordinates normalized to start at 0 for comparison. Orange tracks denote haplotypes transmitted across three generations, and blue tracks denote haplotypes transmitted across two generations. Methylation profiles are shown for Generation 1–3 (Gen1–Gen3). Black bars indicate the position of the centromeric dip region (CDR). **B**, Illustration of Methylation Area Score (MARS) calculation. The figure shows the centromere dip region (CDR) and the surrounding context of PAN027Mat and PAN028Mat. The top gray bar denotes the annotated CDR. Each black point represents the average CpG methylation level of a 1kb window across the region. The shaded blue area highlights the bins used for MARS calculation, defined as the middle 90% of bins after excluding extreme values. Local upper and lower reference baselines are defined as the mean methylation of the top 5% and bottom 5% of bins, respectively. MARS is computed as the summed difference between the upper reference baseline and the observed methylation levels across the shaded region, thereby integrating both the depth and breadth of hypomethylation within the CDR. **C**, Intergenerational variability of CDR-associated metrics quantified by coefficients of variation (CV) for CDR length, mean methylation, and log_2_-transformed methylation area score (MARS). P values were calculated using Wilcoxon rank-sum tests; **** denotes *P* < 0.0001. **D**, Chromosome-wide distribution of log_2_(MARS) across haplotypes. Dark red indicates haplotypes transmitted across three generations (Tr2), orange indicates haplotypes transmitted across two generations (Tr1), and light blue indicates independent haplotypes with no transmission (Tr0). **E–F**, Changes in CDR length (**D**) and MARS (**E**) across cell states. PBMC, peripheral blood mononuclear cells; iPSC, induced pluripotent stem cells; NPC, neural progenitor cells. P values were calculated using a Wilcoxon rank-sum test; **** denotes *P* < 0.0001. **G**, DNA methylation levels within the active HOR (excluding the CDR) of chromosome 1, across PBMCs, iPSCs, and NPCs. **H**, Correlation between changes in hypomethylation of the active HOR (excluding the CDR) and the extent of MARS recovery in chromosome 1. Points are colored by individual (PAN010, PAN011, PAN027, PAN028), and shapes indicate haplotype origin (maternal, circles; paternal, triangles). **I**, DNA methylation profiles across the active HOR of chromosome 1 for different haplotype-resolved assemblies and cell states. Colors indicate cell type (PBMC, dark red; iPSC, light blue; NPC, orange). Symbols denote inheritance status: circles, three-generation transmission (Tr2); triangles, two-generation transmission (Tr1); crosses, independent haplotypes (Tr0). **J**, Expanded view of DNA methylation across the full centromeric satellite region and flanking ±10 Mb of chromosome 1 in PAN011Pat. Colors indicate cell type as in **H.** Dark green circles mark the position of the CDR, and the active HOR is highlighted in red box.

To investigate whether CDR architecture and methylation patterns are maintained during inheritance, as well as during cellular reprogramming and differentiation, we compared CDRs across PBMCs, iPSCs, and NPCs for all haplotypes. Examination of CDR methylation profiles revealed pronounced structural diversity across chromosomes and haplotypes (**Fig. 2A-B; Supplementary Fig. 11**). The shape, depth, and continuity of the methylation valleys varied across haplotypes and chromosomes (**Supplementary Fig. 10**), ranging from simple “U-shaped” depressions (e.g., PAN010 Pat chr1) to multi-valley “W-shaped” profiles (e.g., PAN028 Pat chr2) to broader, undulating troughs (e.g., PAN010 Mat chr3). A small subset looked like two adjacent valleys (e.g., PAN028 Pat chr13), consistent with prior reports describing “dual-dip” architectures^10,13^. Across all haplotypes and chromosomes, the mean CDR length across all haplotypes was ∼305 kb (range 110–660 kb) (**Supplementary Table 7**), with chrY containing the smallest CDR (∼110 kb) and PAN011 Mat chr1 the largest (∼660 kb).

Due to the high diversity of the CDR architecture, single measurements such as CDR length or mean methylation level provide limited information, as they cannot distinguish between different methylation patterns that may represent distinct epigenetic states. The diverse hypomethylation patters likely reflect the heterogeneity among cell populations or intrinsic methylation patterns of the DNA molecules. To capture the full extent of centromeric hypomethylation, we defined a Methylation Area Score (MARS) that quantifies the total integrated hypomethylation across the CDR. For each CDR, we calculated methylation levels in 1kb bins and computed MARS by summing the difference between a local baseline (mean of the top 5% highest-methylated bins) and the observed methylation across the middle 90% of bins, excluding extremes to minimize outlier effects (**Fig. 2B**) (Methods). This metric integrates both the depth and breadth of hypomethylation while remaining robust to local fluctuations. To establish MARS as a stable metric of centromeric methylation architecture, we first examined its behavior across generations. The result shows MARS exhibited substantially lower intergenerational variability, as quantified by the coefficient of variation (CV) across three generations (mean CV: log2(MARS) = 0.01, CDR length = 0.31, mean methylation = 0.07; Wilcoxon rank-sum test: log2(MARS) vs. length, P = 1.2 × 10^−13^; log2(MARS) vs. mean methylation, P = 2.5 × 10^−10^) (**Fig. 2C**). These results indicate that while CDR length and average methylation levels can fluctuate across generations, the global hypomethylated state captured by MARS remains relatively stable.

In contrast to its intergenerational stability, MARS values differed substantially between chromosomes (**Fig. 2D**). For example, chromosome 1 exhibited approximately 1.6-fold higher MARS compared to chromosome 21 (mean log_2_(MARS): 15.77 ± 0.12 vs. 15.11 ± 0.13; P < 0.001, this difference was consistently observed in PBMCs, iPSCs, and NPCs (**Supplementary Fig. 18**). Pairwise comparisons across all chromosomes revealed that 45% (114/253) of chromosome pairs showing significant differences (P < 0.01) (**Fig. 2D**). Notably, chromosomes 4 and Y, previously reported to show reduced CENP-A levels^40,41^, also exhibited lower MARS values, though direct quantification of CENP-A density would be required to establish whether MARS differences systematically reflect kinetochore composition. These observations establish that MARS captures chromosome-specific centromeric methylation architecture that is maintained across inheritance, providing a foundation for examining changes during cell-fate transitions.

### 3 Reprogramming collapses CDR hypomethylation pattern, whereas NPC differentiation partially restores it

Reprogramming is accompanied by extensive genome-wide epigenetic remodeling, yet how the core epigenetic architecture of centromeres responds to this process remains poorly defined. To address this gap, we examined CDR features across PBMCs, iPSCs, and differentiated NPCs across the pedigree to determine whether the epigenetic landscape of centromeres undergoes remodeling during cell-fate transitions.

Having established the stability of MARS across generations, we next asked whether CDR positions and methylation patterns are maintained during reprogramming and differentiation. The center position of each CDR remained highly stable across PBMCs, iPSCs, and NPCs, with positional shifts limited to <50 kb (mean 31.3 kb, SD 33.2 kb) (**Supplementary Figs. 12–16**), comparable to the positional stability observed across generations (mean shift 62.4 kb; **Supplementary Fig. 11**).

Despite this positional stability, methylation patterns changed dramatically during reprogramming. CDRs were substantially narrower in iPSCs compared to PBMCs (PBMCs mean 306.0 kb, SD 102.3 kb; iPSCs mean 217.5 kb, SD 82.3 kb; Wilcoxon P = 2.37 × 10^−19^), and remained comparable in NPCs (mean 220.0 kb, SD 79.5 kb; PBMCs vs. NPCs P = 0.57) (**Fig. 2E**). More strikingly, MARS was markedly reduced in iPSCs, revealing a pronounced attenuation of CDR hypomethylation with an average reduction of ∼49.3% relative to PBMCs (mean log_2_MARS: PBMC 15.50 ± 0.31 vs. iPSC 14.52 ± 0.42; P = 4.36 × 10^−54^) (**Fig. 2F**). During differentiation into NPCs, MARS substantially recovered to ∼91.4% of PBMC levels (mean log_2_MARS: NPCs 15.37 ± 0.35; iPSCs vs. NPCs P = 2.99 × 10^−50^, PBMCs vs. NPCs P = 7.2 × 10^−6^) (**Fig. 2F**). This pattern of CDR attenuation in iPSCs and restoration in NPCs was highly consistent across all haplotypes (**Supplementary Fig. 16**) and chromosomes (**Supplementary Fig. 17**), indicating a centromere-specific regulatory process. These results reveal a previously uncharacterized layer of centromere epigenetic plasticity during cell state transitions, which is concordant with previous reports of CENP-A depletion at centromeres in human iPSCs and embryonic stem cells^25,26^. Importantly, despite the global attenuation of MARS in iPSCs and its restoration in NPCs, the relative chromosome-specific pattern of CDR hypomethylation was largely preserved across cell types (**Supplementary Fig. 18**). Chromosomes 4 and Y consistently exhibited lower MARS compared to other chromosomes in PBMCs, iPSCs, and NPCs.

Beyond the core CDR, other peri- and centromeric satellite families (including active HOR arrays outside the CDR, divergent HORs (dHOR), canonical HORs, and HSat1–3) also exhibited cell-type–specific methylation differences (**Supplementary Figs. 21, 22**). We focused on the CDR-embedded active HOR arrays, which form the functional centromeric core. Using chromosome 1 as an example, chromosome-wide DNA methylation differences among cell types were modest (PBMC: 78.3 ± 0.49%; iPSC: 80.52 ± 1.71%; NPC: 76.17 ± 0.78%). In contrast, active HOR arrays displayed pronounced NPC-specific hypomethylation (PBMC: 87.01 ± 4.10%; iPSC: 88.58 ± 4.24%; NPC: 77.14 ± 7.30%) (**Fig. 2G**). This NPC-specific reduction in methylation was generally consistent across chromosomes and haplotypes (**Supplementary Fig. 19**), with notable exceptions including PAN028 Mat/Pat chr1, PAN028 Mat/Pat chr18, and PAN028 Pat chr20, which remained highly methylated as in iPSCs (**Supplementary Figs. 12, 14, 15, 19**). PAN010 Mat/Pat chr1 exhibited an interesting split pattern in which the region 3’ to the CDR showed typical methylation reduction while the 5’ region remained highly methylated (**Fig. 2I**).

Given that CDR demethylation is correlated with CENP-A binding and that targeted demethylation of active HOR units can promote CENP-A deposition^17^, we asked whether active HOR demethylation is associated with CDR restoration in NPCs. Using PBMCs as a baseline, we quantified (i) the degree of CDR restoration (ΔMARS = [log_2_(MARS_NPC) − log_2_(MARS_iPSC)] / log_2_(MARS_PBMC)) and (ii) the magnitude of active HOR demethylation (ΔHOR_Meth = [iPSC_meth − NPC_meth] / PBMC_meth). Across 23 chromosomes, 7 showed strong correlations (Pearson r > 0.7), 10 showed moderate correlations (0.4 < r < 0.7), and 4 showed weak correlations (0.2 < r < 0.4) (**Supplementary Fig. 20**). For chromosome 1, variation in active HOR demethylation strongly correlated with CDR restoration (r = 0.81, P < 0.001; **Fig. 2H**).

Collectively, these results demonstrate that restoration of CDR methylation architecture during differentiation from iPSCs is associated with coordinated, large-scale demethylation of active HOR arrays. Notably, this demethylation is largely confined to peri- and centromeric regions and does not extend into chromosome arms (**Fig. 2J; Supplementary Figs. 21D, 22D**), supporting a centromere-specific mode of epigenetic remodeling during neural differentiation.

### 4 Chromatin state remodeling at centromeres revealed by Fiber-seq

Fiber-seq leverages the adenine methyltransferase (e.g., Hia5) to deposit m6A on accessible DNA, whereas DNA wrapped around nucleosomes or occluded by protein complexes is protected from m6A modification^38,42^. Importantly, this process does not interfere with endogenous CpG (5mC) methylation detection. As a result, each long-read fiber simultaneously encodes multiple layers of chromatin information, including CpG methylation, nucleosome footprints, protein-bound footprints (FIREs, Fiber-seq Inferred Regulatory Elements), and linker regions. Across all samples, the overall nucleosome footprint length was 133 bp (**Supplementary Fig. 23**), consistent with estimates in previous Fiber-seq studies^42^.

We first examined FIRE enrichment across active HOR arrays (Methods). As expected, FIRE signals formed prominent peaks at CDRs (**Fig. 3A; Supplementary Figs. 24, 25**), consistent with the localization of centromere-associated protein complexes, including CENP-C, CENP-N, and other constitutive centromere-associated network (CCAN) components^43,44^. To quantify protein occupancy, we calculated a FIRE Area Score (FAS; Methods) for each CDR. Differentiation of iPSCs into NPCs was associated with significant ∼13% increase of FAS (mean log_2_(FAS) = 12.14 ± 0.41 in iPSCs vs. 12.32 ± 0.41 in NPCs; Wilcoxon test, P = 6.93 × 10^−6^) (**Fig. 3B**), which corroborated the partial restoration of CDR hypo-methylation during NPC differentiation. These results define profound functional epigenetic reprogramming signatures of lineage commitment at the centromere: reduced DNA methylation, enhanced protein occupancy, and chromatin remodeling.

**Figure 3.**
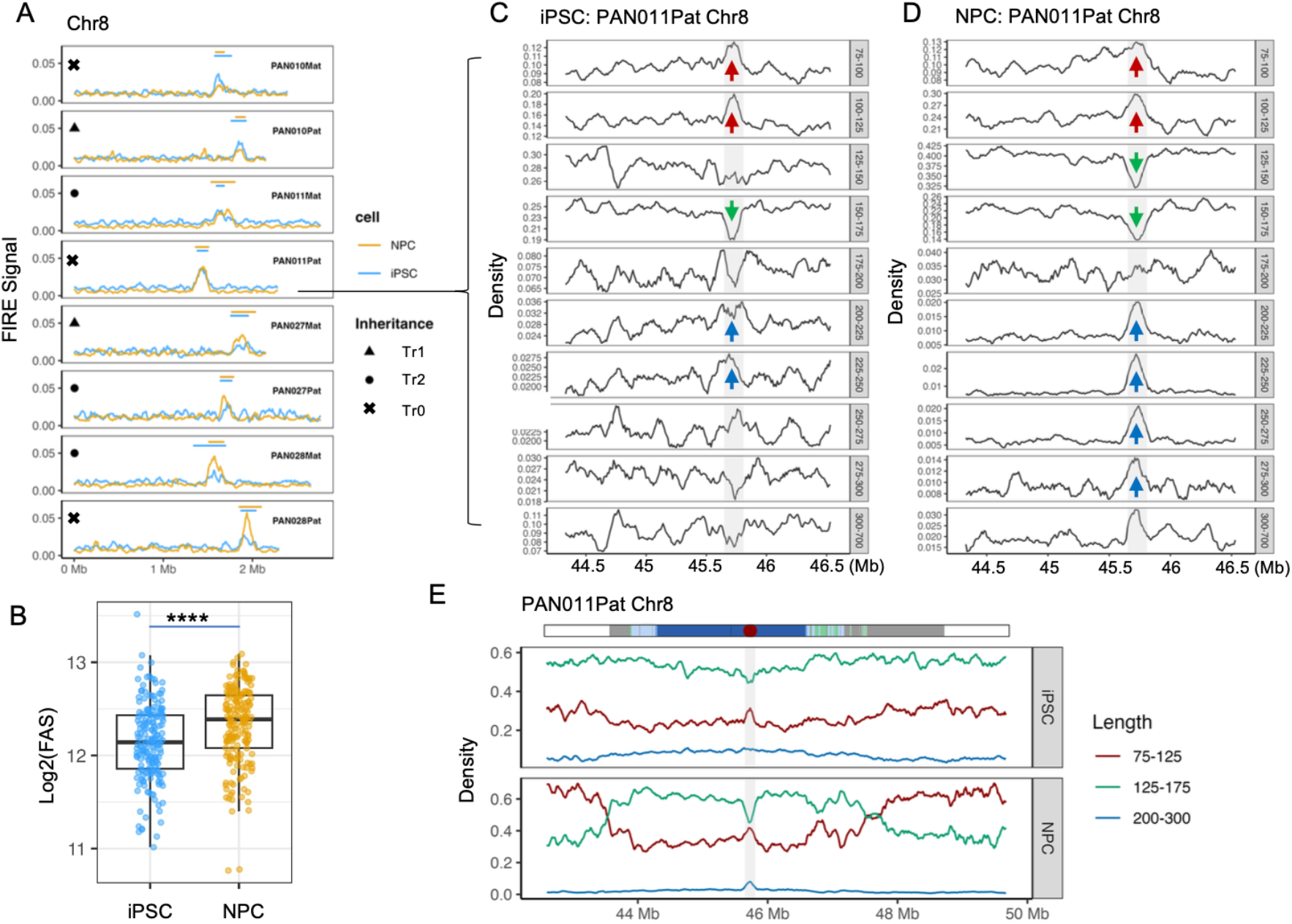
Reprogramming alters nucleosome organization within centromeric dip regions, with partial restoration upon NPC differentiation. **A**, Fiber-seq–derived FIRE (Fiber-seq Inferred Regulatory Element) signals across the active HOR of chromosome 8 for multiple haplotype-resolved assemblies. FIRE profiles are shown for induced pluripotent stem cells (iPSCs, blue) and neural progenitor cells (NPCs, orange). Symbols denote inheritance status of each haplotype: circles indicate haplotypes transmitted across three generations (Tr2), triangles indicate haplotypes transmitted across two generations (Tr1), and crosses represent independent haplotypes (Tr0). **B**, Comparison of CDR FIRE area scores (FAS) between iPSCs and NPCs across all chromosomes. Each point represents a haplotype. P values were calculated using a Wilcoxon rank-sum test; **** denotes *P* < 0.0001. **C–D**, Length-resolved nucleosome footprint density profiles across active HOR of chromosome 8 for PAN011Pat in iPSCs (**C**) and NPCs (**D**). Each subpanel corresponds to a specific nucleosome footprint length range (indicated on the right). Red arrows mark regions showing increased enrichment of short nucleosome footprints (75–125 bp), green arrows indicate depletion of canonical-sized nucleosomes (125–175 bp), and blue arrows highlight enrichment of larger nucleosome footprints (200–300 bp). Grey shaded region indicating the position of the centromeric dip region (CDR). **E**, Expended view of distribution of nucleosome footprint densities across the full centromeric satellite (cenSat) region of PAN011Pat chromosome 8 and flanking genomic regions. The cenSat annotation track is shown at the top, with the red dot and grey shaded region indicating the position of the centromeric dip region (CDR). Profiles are shown separately for iPSCs (top) and NPCs (bottom). Red traces denote short nucleosome footprints (75–125 bp), green traces denote canonical-sized nucleosomes (125–175 bp), and blue traces denote larger nucleosome footprints (200–300 bp).

CENP-A–containing nucleosomes are known to wrap ∼120 bp of DNA^45,46^, shorter than canonical H3 nucleosomes (∼147 bp). Fiber-seq enables direct, single-molecule measurement of nucleosome footprint lengths^15,47^ (**Methods**), allowing us to test whether distinct α-satellite subdomains harbor structurally distinct nucleosomes. We extracted all nucleosome footprints between 75 and 700 bp from phased Fiber-seq reads and examined length distributions across active HOR arrays. As expected, shorter footprints (75–125 bp) were markedly enriched within CDRs, accompanied by a depletion of nucleosomes of canonical size (125–175 bp). Strikingly, we also observed a pronounced enrichment of larger footprints (200–300 bp) within CDRs (**Fig. 3D**). These larger footprints are consistent with recent cryo-electron tomography observations showing that, in addition to the CCAN complex associated with individual CENP-A nucleosomes, centromeric regions contain larger globular protein assemblies^48^. Notably, this feature was more prominent in NPCs than in iPSCs (**Fig. 3C**) and was consistently observed across haplotypes (**Supplementary Fig. 26**), further supporting increased protein occupancy and chromatin reorganization at CDRs in differentiated cells.

Extending this analysis to the entire cenSat domain revealed broad remodeling of nucleosome architecture during differentiation. In iPSCs, both cenSat and flanking regions were dominated by nucleosomes of canonical size (∼60%). Upon differentiation into NPCs, canonical nucleosomes were selectively depleted at CDRs, accompanied by an increase in footprints of shorter nucleosomes, whereas pericentromeric satellite subdomains exhibited substantial increase of shorter nucleosome occupancy (from ∼40% to ∼60%), a pattern consistently observed across haplotypes (**Supplementary Fig. 27**). Together, these results demonstrate that centromeric chromatin—particularly within CDRs—undergoes extensive, differentiation-associated remodeling of nucleosome architecture, revealing a structural layer of regulation that operates alongside, but is not fully reflected by, DNA methylation changes.

### 5 X-inactivation leaves broad scale epigenetic memory in chromosome arms but not in the centromere

Chromosome X inactivation (Xi) is a pervasive feature of female somatic cells, whereas reprogramming to induced pluripotent stem cells (iPSCs) frequently induces heterogeneous erosion of X inactivation (Xe)^28^. Both processes involve extensive epigenomic remodeling, including large-scale changes in DNA methylation^27,29^. We therefore asked whether X-linked epigenetic reprogramming extends into centromeric chromatin.

A central regulator of X inactivation is X inactive specific transcript (XIST), a long noncoding RNA expressed exclusively from the inactive X chromosome^49^. Accordingly, XIST expression and promoter methylation serve as robust indicators of X-chromosome status: the Xi allele is characterized by hypomethylation at the XIST promoter, whereas the active X (Xa) remains hypermethylated. Haplotype-resolved CpG methylation profiling unambiguously identified the Xi allele in PBMCs from all three female individuals (**Fig. 4A–C**). Specifically, the inactive haplotypes corresponded to PAN010 Pat, PAN027 Mat, and PAN028 Pat, each displaying large allelic methylation differences at the XIST promoter (Δmeth 0.33–0.71) (**Fig. 4D–F**). Consistent with the mosaic Turner karyotype of PAN028 (45,X/46,XX), PBMCs from this individual exhibited intermediate promoter methylation on both alleles, resulting in a reduced maternal–paternal difference compared with PAN010 and PAN027.

**Figure 4.**
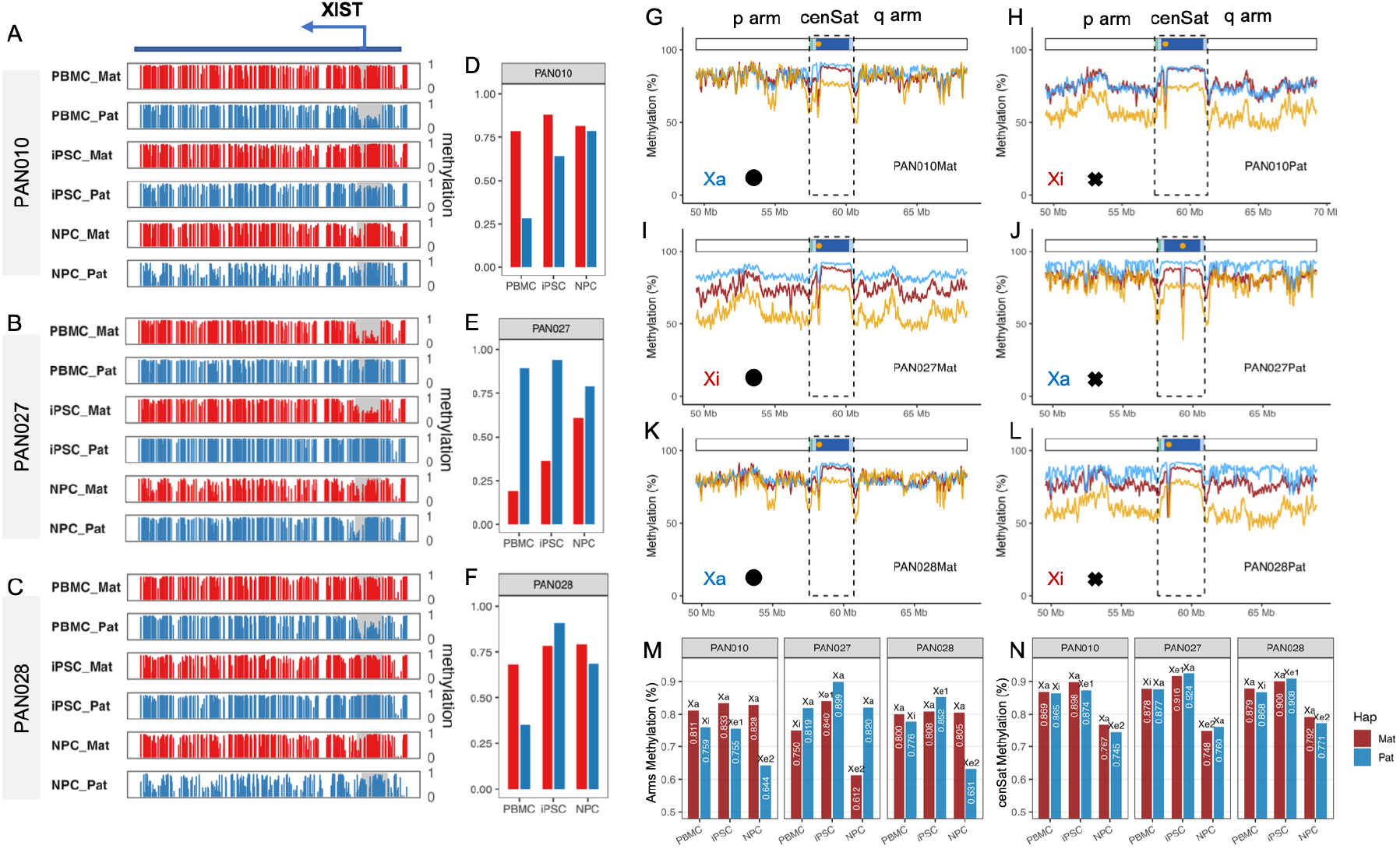
X-chromosome inactivation–associated DNA methylation patterns across chromosome arms and centromeres. **A–C**, Haplotype-resolved CpG methylation profiles across the *XIST* locus in PAN010 (**A**), PAN027 (**B**), and PAN028 (**C**). Maternal (red) and paternal (blue) alleles are shown for peripheral blood mononuclear cells (PBMCs), induced pluripotent stem cells (iPSCs), and neural progenitor cells (NPCs). The grey shaded region indicates the *XIST* promoter, whose mean methylation levels are summarized in **D–F. D–F**, Mean CpG methylation levels at the *XIST* promoter for each haplotype across PBMCs, iPSCs, and NPCs in PAN010 (**D**), PAN027 (**E**), and PAN028 (**F**). Promoter methylation states are used to assign X-chromosome inactivation status. **G–L**, Allele-resolved CpG methylation profiles across the chromosome X centromeric satellite domain (cenSat) and ± 8Mb flanking regions for each haplotype. The upper annotation track denotes cenSat substructure, and orange dots mark the positions of centromeric dip regions (CDRs). In each figure, the line colors denote different cells (PBMCs, dark red; iPSCs, sky blue; NPCs, orange). Symbols denote cenSat inheritance status: black dots indicate transmitted across three generations, and crosses represent independent haplotypes. **M**, Average CpG methylation levels across chromosome X arms for different X-chromosome states. **N**, Average CpG methylation levels across centromeric satellite regions for different X-chromosome states. Xa, active X chromosome; Xi, inactive X chromosome; Xe, eroded X chromosome; Xe1, X-chromosome erosion observed in iPSCs; Xe2, X-chromosome erosion observed in NPCs; Mat, maternal allele; Pat, paternal allele.

Reprogramming and differentiation produced distinct trajectories of X-inactivation erosion across individuals^29^. In PAN010 and PAN028, the Xi allele gained substantial XIST promoter methylation during reprogramming, consistent with early erosion of XCI (PAN010 Δmeth reduced from 0.51 to 0.24; PAN028 from 0.33 to –0.12), and this hypermethylated promoter state was largely maintained upon differentiation to NPCs (**Fig. 4D, F**). In contrast, PAN027 showed limited erosion in iPSCs but continued progressive erosion during neural differentiation, with Δmeth decreasing from 0.71 (PBMC) to 0.58 (iPSC) and further to 0.18 (NPC) (**Fig. 4E**).

Beyond the XIST locus, Xi status was reflected in chromosome-wide methylation patterns. In PBMCs, the inactive X was globally hypomethylated relative to Xa (mean chromosome arm Δmeth Xa–Xi = 4.85%). During erosion in iPSCs, despite promoter hypermethylation at XIST, global methylation differences between Xa and Xe remained comparable (mean arm Δmeth Xa– Xe_1_ = 6.00%). In contrast, NPCs showed a pronounced global hypomethylation of the eroded X chromosome (mean arm Δmeth Xa–Xe_2_ = 18.88%), consistent with prior observations that late Xe accompanied by global X hypomethylation^27^. Here, we resolved this pattern at haplotype and telomere to telomere resolution (**Fig. 4 H, I, L, M**).

We next asked whether differences in X-chromosome status (Xa, Xi, Xe) extend into the centromere. Similar to autosomes, centromere epigenomic structure remained highly stable across all X states, with only minor positional shifts (<30 kb) across generations and cell types (**Fig. 4G–L; Supplementary Fig. 28**). Strikingly, methylation differences within peri/centromeric region between X alleles were minimal in stark contrast to chromosome arms (mean active HOR allelic Δmeth: 0.55% in PBMCs, 1.35% in iPSCs, and 1.85% in NPCs, compared to arm’s allelic Δmeth Wilcoxon signed-rank test p = 0.004). This result indicates that centromere methylation is largely independent of Xa, Xi, or Xe status. In addition, the NPC-associated global hypomethylation of active HOR arrays was comparable between Xa and Xe chromosomes (mean Δmeth: 13.42% in Xa vs. 14.38% in Xe, Wilcoxon signed-rank test p = 0.42) (**Fig. 4N; Supplementary Table 9**). Notably, late-stage Xe induced stronger hypomethylation across chromosome arms (Δmeth 18.88%) than observed within centromeric satellite DNA during differentiation (Δmeth 13.90%). This result suggests that XCI erosion–associated hypomethylation and differentiation-associated centromeric hypomethylation represent mechanistically distinct processes, potentially governed by different regulatory constraints.

### 6 Centromeres accumulate the highest burden of reprogramming-associated de novo mutations

Genomic stability is a critical consideration for the safety of iPSCs in disease modeling and potential therapeutic applications. To quantify de novo variants arising during reprogramming, we leveraged haplotype-resolved assemblies and assessed newly acquired variants in iPSCs and NPCs relative to donor-matched PBMCs.

Phased PacBio HiFi reads were aligned to their corresponding maternal or paternal assemblies, and de novo variants were identified using two complementary pipelines, with stringent filtering to retain high-confidence events (Methods). PAN028 showed markedly reduced concordance between iPSC and NPC variant sets and was excluded from downstream enrichment and mutational spectrum analyses (**Supplementary Note 1**).

Because NPCs were derived from iPSCs over a short differentiation interval, variants emerging in iPSCs were expected to be inherited by NPCs, with NPCs potentially accumulating slightly more variants. Consistent with this expectation, iPSCs carried an average of 685 (628–771) de novo variants per haplotype assembly, including 657 SNPs (596–746) and 28 short indels (24– 33). NPCs exhibited a modestly higher number, with 725 (641–798) variants, comprising 694 SNPs (611–770) and 31 indels (24–39) (**Fig. 5A; Supplementary Table 10**). Cross-validation between iPSCs and NPCs revealed strong concordance: 92.96% (89.23–95.19%) of iPSC variants were also detected in NPCs, including 93.39% (89.68–96.41%) of SNPs and 84.02% (72.73–95.83%) of indels (**Fig. 5B**). These results show that most reprogramming-associated mutations are faithfully inherited during early neural differentiation. Notably, the two independent PAN027 iPSC clones carried highly similar numbers of de novo variants (maternal haplotype: 660 in Clo1 vs. 656 in Clo2; paternal haplotype: 669 in Clo1 vs. 657 in Clo2) (**Supplementary Table 10**), yet none of these variants were shared between the clones, demonstrating that these mutations arose independently in each clone rather than originating from a common parental cell.

**Figure 5.**
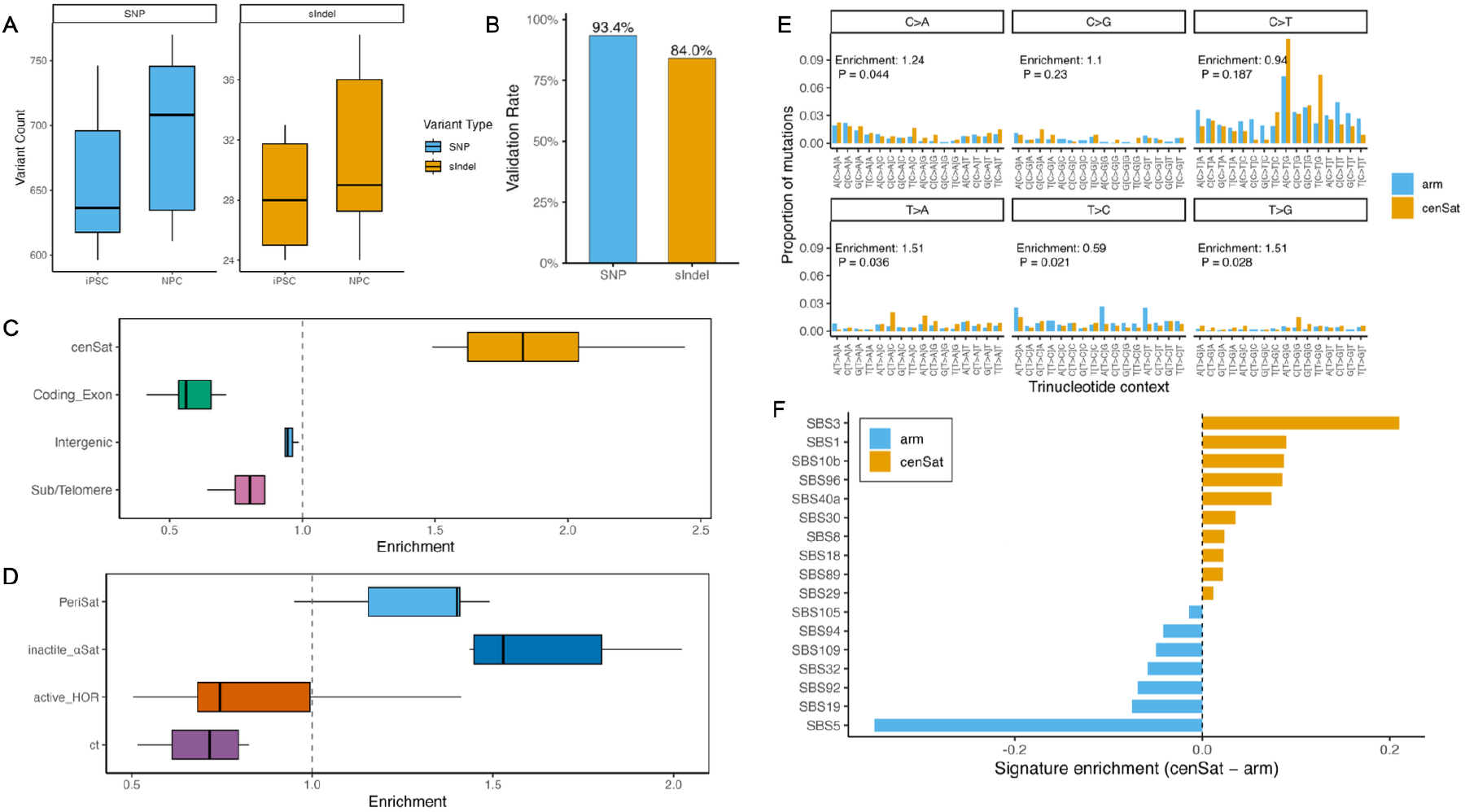
De novo variant burden and mutational spectra across centromeric subdomains. **A**, Counts of de novo single-nucleotide variants (SNPs) and short insertions/deletions (sIndels) detected in induced pluripotent stem cells (iPSCs) and neural progenitor cells (NPCs) across haplotype-resolved assemblies. **B**, Validation of iPSC-derived de novo variants in NPCs. Bars indicate the fraction of variants identified in iPSCs that are also detected in NPCs, shown separately for SNPs and sIndels. **C**, Enrichment of de novo variants across major genomic compartments, normalized by the genomic span of each compartment. Shown are centromeric satellites (cenSat), coding exons, intergenic regions, and subtelomeric regions. **D**, Enrichment of de novo variants across functional centromeric subdomains, including active higher-order repeats (active_HOR), inactive α-satellite, pericentromeric satellite regions (periSat), and centromere transition regions (ct). **E**, Trinucleotide mutational spectra (SBS96) of de novo single-nucleotide variants in chromosome arm regions and cenSat regions. Mutation frequencies are shown for each of the six base substitution classes, stratified by trinucleotide context. Enrichment values and corresponding *P* values indicate differences between cenSat and chromosome arms. **F**, Differential contributions of COSMIC single base substitution signatures to mutational spectra in cenSat regions relative to chromosome arms. Bars represent the difference in signature exposure (cenSat − arm), with positive values indicating cenSat enrichment and negative values indicating arm enrichment.

To determine whether de novo mutations are evenly distributed across genomic compartments, we quantified enrichment in four major regions: (1) centromeric satellites (cenSat), (2) subtelomeric regions (within 1 Mb of chromosome ends), (3) coding exons (excluding those overlapping centromere or subtelomere), and (4) the remainder of the genome. Enrichment was computed as: (mutations in region / total mutations) / (region size / assembly size).

cenSat regions exhibited a markedly higher mutation burden than other compartments, with an enrichment of 1.88 (1.49–2.44), whereas coding exons showed strong depletion, 0.58 (0.41–0.71). The mutation rate in cenSat was therefore 3.31-fold (2.21–3.89) higher than in coding exons (**Fig. 5C**).

We next dissected mutational patterns within distinct centromere subdomains. Based on sequence composition and functional annotation, we categorized centromeric repeats into four classes: (1) active_HOR—the kinetochore-forming higher-order repeat array; (2) inactive α-satellite (“shell α-satellite”), including dHOR, HOR, mixedAlpha, monomeric α-satellite, and other cenSat repeats; (3) pericentromeric satellites, including HSat2, HSat3, HSat1A, HSat1B, gSat, and bSat; (4) the centromere transition (ct) region, consisting predominantly of non-repetitive sequence adjacent to the α-satellite arrays.

Mutation rates differed substantially across these domains. active_HOR and ct regions exhibited strong depletion of de novo variants relative to all other centromeric and pericentromeric satellites (**Fig. 5D**), suggesting that the core kinetochore-forming domain and surrounding transition region may be subject to enhanced protection from mutational processes. In contrast, inactive α-satellite and pericentromeric satellites accumulated significantly more variants.

We further examined the mutational spectrum within centromeric satellite regions. Compared with chromosome arms, cenSat regions exhibited a pronounced shift in mutational patterns, with a cosine similarity of 0.84 between arm and cenSat. Trinucleotide-based analysis (**Methods**) revealed that cenSat regions were significantly enriched for T>A and T>G substitutions (1.51-fold, P = 0.036 and P = 0.028, respectively), while T>C substitutions were significantly depleted (0.59-fold, P = 0.021) (**Fig. 5E; Supplementary Fig. 29**).

To investigate the underlying mutational processes, we decomposed the 96-channel mutational profiles (SBS96) using COSMIC^50^ v3.5 signatures. Chromosome arms were dominated by the clock-like mutational signature SBS5 (NNLS exposure = 0.35 in arms, 0.00 in cenSat), whereas cenSat regions were enriched for signatures associated with alternative mutational processes, including the replication- and repair–associated signature SBS3 (0.00 in arms, 0.21 in cenSat) and the CpG deamination–associated signature SBS1 (0.15 in arms, 0.24 in cenSat) (**Fig. 5F, Supplementary table 11**). These results are consistent with distinct DNA damage and repair dynamics operating in highly repetitive centromeric heterochromatin.

Collectively, these findings reveal that de novo mutations are preferentially concentrated within highly repetitive centromeric satellites and that the mutational processes acting on centromeres differ markedly from those in the rest of the genome. These findings indicate that centromeric regions represent a hotspot for reprogramming-associated mutagenesis, underscoring the importance of monitoring satellite stability in iPSC-based applications.

## Discussion

Centromeres are paradoxical genomic domains: they are essential for chromosome segregation yet embedded within highly repetitive DNA that has historically resisted genetic and epigenomic interrogation. By integrating fully phased telomere-to-telomere assemblies with long-read epigenomic profiling across a three-generation pedigree and matched reprogramming and differentiation states, our study resolves this paradox and reveals fundamental principles governing human centromere organization.

### Centromere position is anchored, while epigenomic capacity is tunable

To integrate our observations across inheritance, reprogramming, and differentiation, we summarize centromeric chromatin remodeling in a conceptual model (**Fig. 6**). This model captures two profound features of centromere organization: a highly stable positional identity and a dynamically regulated epigenomic state. In iPSCs, CDRs are associated with attenuated hypomethylation, reduced protein occupancy, and a relative predominance of canonical-size nucleosomes. Upon differentiation into NPCs, CDRs undergo coordinated remodeling characterized by increased protein-bound footprints, enrichment of short CENP-A–like nucleosomes, and the emergence of extended protected regions. Notably, these changes are spatially restricted to centromeric satellite domains and do not extend into flanking chromosome arms, indicating a centromere-specific chromatin remodeling program during lineage commitment.

**Figure 6.**
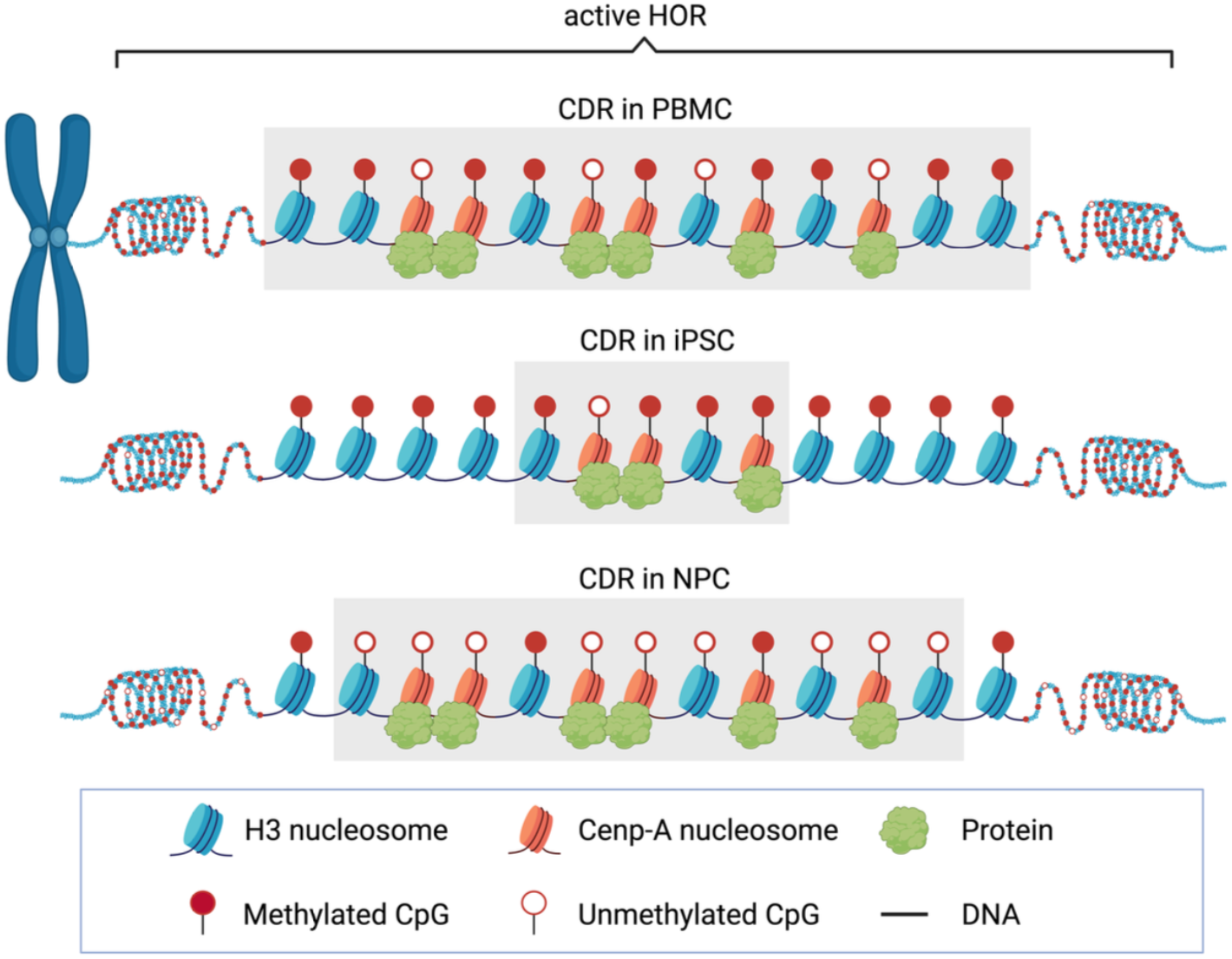
Conceptual model of centromeric chromatin remodeling during cell-fate transitions. Schematic summary of centromeric chromatin dynamics across cell states. Reprogramming to pluripotency is associated with attenuation of centromeric protein occupancy and disruption of specialized nucleosome organization within centromeric dip regions (CDRs). Upon neural differentiation, CDR-associated chromatin features are partially restored, including increased protein binding and the emergence of heterogeneous nucleosome architectures. Throughout these transitions, the positional identity of centromeres is preserved, indicating a separation between centromere localization and epigenomic remodeling.

The remarkable stability of CDR positions across inheritance, reprogramming, and differentiation is consistent with the unique inheritance properties of centromeric CENP-A chromatin. CENP-A nucleosomes are long-lived and propagate centromere identity through a self-templated mechanism, in which inherited CENP-A specifies the site of new deposition after DNA replication^43,51^. Our allele-resolved analyses provide direct evidence that this mechanism anchors centromere position with exceptional fidelity, even during extensive epigenomic resetting.

In contrast, centromere epigenome is highly plastic. We observe a pronounced attenuation of CDR hypomethylation in iPSCs, followed by partial restoration during NPC differentiation, without major shift in centromere position. Thus, reprogramming and differentiation modulate centromeric chromatin state rather than positional identity. Such modulation is compatible with known features of pluripotent cell cycles^52^ and reported reductions of centromere-localized CENP-A in pluripotent states^25,26^, which may constrain centromeric chromatin status during reprogramming while leaving positional memory intact.

Prior work has suggested that functional centromeres operate under quantitative constraints on CENP-A–containing chromatin^40^. Our results extend this concept by revealing an epigenomic correlate of such regulation. Despite fluctuation in fine-scale methylation patterns, the integrated hypomethylated capacity of CDRs, measured by MARS, is strongly conserved across inheritance and reversibly tuned across cell states. Together, these findings support a model in which centromere position is stably inherited, while centromeric epigenomic capacity is quantitatively constrained yet developmentally plastic.

### Chromatin remodeling at centromeres during lineage commitment

Centromere epigenetic remodeling during differentiation extends beyond DNA methylation. The enrichment of protein-bound footprints and non-canonical nucleosome footprints within CDRs in NPCs indicates increased protein occupancy and altered chromatin topology upon lineage commitment. These features are consistent with re-establishment of kinetochore-proximal chromatin organization^45,53^ following the epigenetic disruption induced by reprogramming.

Selective depletion of canonical-length nucleosomes and enrichment of short and extended protected regions within CDRs highlight the structural diversity of centromeric chromatin. Such heterogeneity likely facilitates accommodation of specialized nucleosomes and large protein assemblies at the kinetochore, underscoring the importance of single-molecule approaches for resolving centromeric chromatin architecture.

### Insulation of centromeres from chromosome-wide epigenetic programs

Despite profound methylation differences across chromosome arms associated with X-chromosome inactivation and erosion, centromeric methylation patterns and positions remain remarkably stable. This insulation indicates that centromeres are governed by regulatory mechanisms distinct from chromosome-wide epigenetic programs.

At the same time, haplotypes that have experienced X inactivation in vivo exhibit greater epigenetic variability during differentiation, revealing a form of long-term epigenetic memory. Thus, while centromeres themselves are buffered against X-linked regulatory states, broader chromosomal history can still shape epigenomic responsiveness elsewhere along the chromosome.

### Centromeric satellites as hotspots of reprogramming-associated mutagenesis

Haplotype-resolved variant analysis demonstrates that centromeric satellite DNA accumulates the highest burden of de novo mutations during reprogramming, exceeding that observed in coding regions by approximately threefold. Notably, mutation rates are not uniform across centromeric domains: active HOR arrays and centromere transition regions are strongly depleted for mutations, whereas inactive α-satellite and pericentromeric satellites accumulate substantially more variants.

This stratification suggests differential protection across centromeric domains, potentially mediated by chromatin state, protein occupancy, or replication timing. The depletion of mutations within active HORs further supports their functional importance and implies selective constraints that preserve kinetochore integrity. These findings highlight centromeric satellites as sensitive indicators of genome instability that are largely inaccessible to short-read analyses, with implications for iPSC-based applications.

### Developmental vulnerability during centromere epigenetic restoration

Although CDR methylation patterns were largely restored during NPC differentiation, we observed a small number of outlier chromosomes with incomplete recovery, disproportionately affecting PAN028 (Mat/Pat Chr1, Mat/Pat Chr18, and Pat Chr20), the only individual with documented mosaic Turner karyotype. Notably, pluripotent states are known to exhibit elevated chromosomal instability relative to differentiated cells^54^, reflecting transient vulnerabilities during epigenomic resetting.

While PAN028 mosaicism predominantly involves the X chromosome, the enrichment of CDR recovery outliers raises the possibility that early chromosomal or epigenetic perturbations may influence the fidelity of centromeric epigenetic re-establishment more broadly. Although speculative, these observations suggest that incomplete restoration of centromere-associated epigenomic features during pluripotency exit may represent a general vulnerability with potential relevance to developmental disorders and disease.

### Implications and future directions

Together, our study reframes centromeres as genomic domains defined by the coexistence of architectural stability and epigenetic flexibility. This duality likely underlies the evolutionary success of centromeres, allowing them to preserve essential functions while remaining responsive to developmental cues.

Important questions remain. How are centromeric epigenomic states re-established during differentiation, and which molecular pathways coordinate this process? Do elevated mutation rates in centromeric satellites contribute to long-term instability in iPSC-derived cells? Addressing these questions will require integrating telomere-to-telomere assemblies with temporal and perturbation-based epigenomic profiling and functional assays of centromere activity.

By providing a haplotype-resolved, telomere-to-telomere framework spanning inheritance, reprogramming, and differentiation, our study establishes a foundation for dissecting how one of the most essential—and elusive—genomic structures balances stability and change.

## Author Contributions

Ting Wang, Karen H. Miga and Shihua Dong conceived the study. Shihua Dong, Monika Cechova performed data analysis. Monika Cechova conducted genome assembly. Hailey Loucks developed cenSat annotation workflow. Xiaoyun Xing and Heather Schmidt performed sequencing and experimental work. Amber Neilson, Selvamani Vijayalingam, Monica Sentmanat, and Xiaoxia Cui carried out reprogramming and differentiation experiments. Shihua Dong wrote the original draft of the manuscript. Ting Wang, Monika Cechova, Xiaoxia Cui, Ming Hu and Karen H. Miga reviewed and edited the manuscript. Juan Macias, Tianjie Liu, Zheng Dong, Benpeng Miao, Wenjin Zhang, Chad Tomlinson and Eddie Belter provided additional data analysis support. All authors reviewed and approved the final manuscript.

## Acknowledgements

We thank members of the Wang laboratory and the Miga laboratory for helpful discussions. We acknowledge the Genome Technology Access Center and High Performance Computing facilities at Washington University in St. Louis for sequencing and computational support. We thank colleagues in the Telomere-to-Telomere and Human Pangenome Reference Consortium communities for making reference resources publicly available. This work was supported by National Institutes of Health (NIH) grants U41HG010972, U41HG010971, U24HG012070, R01HG007175, R35HG011922.

## Competing Interests

The authors declare no competing interests.

## Declaration of generative AI and AI-assisted technologies in the writing process

During the preparation of this manuscript, the authors used ChatGPT to assist with specific programming-related tasks and to improve the clarity and organization of the text. All content generated with the assistance of this tool was carefully reviewed, edited, and verified by the authors. The authors take full responsibility for the integrity and accuracy of the final manuscript.

## Notes

### Competing Interest Statement

The authors have declared no competing interest.

