## Supplemental Methods for "Fully T2T pedigree assemblies reveal genetic stability and epigenetic plasticity of human centromeres across inheritance and cell-fate transitions"

**Supplementary Materials**

**Supplementary Methods**

1 **iPSC reprogramming**

Peripheral blood mononuclear cells (PBMCs) were obtained from four pedigree members (PAN010, PAN011, PAN027, PAN028). PBMCs were reprogrammed into induced pluripotent stem cells (iPSCs) using Sendai virus vectors (CytoTune™-iPS 2.0 Sendai Reprogramming Kit; Thermo Fisher Scientific) delivering the four Yamanaka factors OCT4, SOX2, KLF4, and c-MYC. Transductions were performed following the manufacturer’s protocol.

All iPSC lines were derived from single-cell clones. For PAN027, two independent clonal iPSC lines were established. For PAN028, PBMCs exhibited mosaicism; the selected iPSC clone used in this study originated from a 46,XX cell.

Karyotype analysis was performed by the Department of Laboratory and Genomic Medicine, Washington University in St. Louis, using standard GTW (G-banding with trypsin and Wright staining) cytogenetic methods. Briefly, iPSCs were treated with a hypotonic solution, fixed, and stained for chromosome analysis. A total of 20 metaphase spreads were examined per line, among which 5 spreads were analyzed and 3 were fully karyotyped at 450-band resolution.

2 **Neural progenitor cell (NPC) differentiation protocol**

Human induced pluripotent stem cells (iPSCs) were differentiated into neural progenitor cells (NPCs) using an embryoid body–based neural induction strategy with dual SMAD inhibition. Neural induction was performed using the STEMdiff™ SMADi Neural Induction Kit (Catalog #08581, STEMCELL Technologies), following the manufacturer’s instructions as described in the Technical Manual for Generation and Culture of Neural Progenitor Cells Using the STEMdiff™ Neural System (Document #10000005588). Briefly, iPSCs were aggregated to form embryoid bodies and subjected to neural induction under SMAD pathway inhibition. After neural commitment, NPCs were maintained and expanded in STEMdiff™ Neural Progenitor Medium (Catalog #05833, STEMCELL Technologies) according to the manufacturer’s protocol. NPC cultures were passaged and expanded under standard conditions prior to downstream molecular and epigenomic analyses.

NPC identity was validated by immunofluorescence staining for canonical neural progenitor markers PAX6 and Nestin, with DAPI used for nuclear counterstaining. For immunofluorescence analysis, NPCs were fixed with 4% paraformaldehyde, permeabilized with Triton X-100, and blocked in serum-containing buffer. Cells were incubated with primary antibodies against PAX6 and Nestin, followed by fluorophore-conjugated secondary antibodies and DAPI staining.

3 **PacBio HiFi sequencing**

PBMCs or cultured iPSCs and NPCs were lysed, and high–molecular weight genomic DNA was extracted using standard protocols. Genomic DNA was sheared targeting a modal fragment length of ~20 kb. SMRTbell libraries were prepared following the PacBio protocol using SMRTbell Prep Kit 3.0. Libraries were size-selected on a Sage PippinHT system using a 0.75% agarose cassette (15–20 kb High Pass, 75E configuration) with a minimum fragment size cutoff of 15 kb. Size-selected libraries underwent annealing, polymerase binding, and cleanup (ABC) using PacBio Revio polymerase kits (SPRQ polymerase kit or Revio polymerase kit). Sequencing was performed on the PacBio Revio platform using Revio SMRT Cells and sequencing plates according to the manufacturer’s specifications.

4 **Oxford Nanopore sequencing**

Oxford Nanopore sequencing libraries were prepared using the Ligation Sequencing Kit LSK114. Libraries were sequenced on the PromethION 24 platform using R10.4.1 flow cells. Data acquisition was performed using MinKNOW version 23.07. Basecalling was conducted using the SUP model with Dorado (version 7.4.1). Oxford Nanopore long-read data were used to complement PacBio HiFi sequencing for genome assembly validation and structural characterization.

5 **Fiber-seq library preparation from iPSCs and NPCs**

For induced pluripotent stem cells (iPSCs) and neural progenitor cells (NPCs), Fiber-seq libraries were prepared following established protocols with minor modifications. Briefly, nuclei were isolated from cultured cells under conditions that preserve native chromatin structure. Isolated nuclei were treated in situ with the adenine methyltransferase Hia5 in the presence of S-adenosylmethionine (SAM) as the methyl donor, enabling deposition of N6-methyladenine (m6A) at accessible DNA regions.

Following Hia5 treatment, nuclei were lysed and high–molecular weight genomic DNA was purified. Fiber-seq libraries were then constructed using PacBio HiFi library preparation protocols, identical to those used for standard HiFi sequencing, and sequenced on the PacBio Sequel II or Revio platforms.

This approach enables simultaneous recovery of long-read DNA sequence, endogenous CpG methylation (5mC), and chromatin accessibility features—including nucleosome footprints and protein-bound regions—from individual DNA molecules.

6 **Sequencing overview**

PBMC samples were sequenced using standard long-read genomic DNA libraries (PacBio HiFi and/or ONT), whereas iPSC and NPC samples were sequenced using Fiber-seq libraries generated from Hia5-treated nuclei. Sequencing depth and coverage statistics for all samples are summarized in Supplementary Table 5.

7 **Genome assemblies and haplotype framework**

Fully phased telomere-to-telomere (T2T) diploid genome assemblies for all individuals were generated and validated in a companion study by our collaborators^1^. Briefly, these assemblies represent complete haplotype-resolved reconstructions of all autosomes and sex chromosomes, including fully resolved centromeric satellite arrays. Assembly construction, phasing, validation, and quality control procedures are reported in that study.

For the analyses presented here, we used the finalized haplotype-resolved assemblies as fixed genomic references. Regions flagged as low-confidence or assembly gaps by Flagger^2^ and NucFlag^3^ in the companion study were excluded from all downstream analyses and treated as masked intervals. Assembly coordinates and haplotype identifiers were harmonized across individuals to enable consistent cross-generation and cross-cell-state comparisons.

8 **Identification of crossover (CO) events**

Meiotic crossover (CO) events were identified using a sequence-based, haplotype-resolved approach that does not rely on public reference genomes. For each offspring haplotype, the corresponding haplotype assembly was treated as the reference and compared against both parental haplotype assemblies using dipcall^4^. For example, PAN027Mat was compared independently to PAN010Mat and PAN010Pat.

Genetic variants were called between the offspring and parental assemblies, and genotype similarity was evaluated in sliding genomic windows along each chromosome. Within each window, similarity to each parental haplotype was quantified based on concordant variant states, enabling identification of large inherited haplotype blocks and approximate recombination boundaries as transitions in parental similarity.

To refine crossover breakpoints at higher resolution, homologous sequences flanking each candidate recombination interval were extracted from the parental and offspring assemblies and subjected to local sequence realignment. This step reduced ambiguity introduced by genome-wide alignment noise and repetitive sequence context, allowing precise localization of recombination intervals based on informative variant patterns at the block boundaries.

Using this strategy, recombination intervals were resolved at base-pair to kilobase resolution across autosomes and sex chromosomes, including regions proximal to centromeric and acrocentric domains.

9 **Phasing of PacBio HiFi and Fiber-seq reads**

PacBio HiFi reads, including Fiber-seq datasets, were independently aligned to the maternal and paternal haplotype assemblies using pbmm2 (v1.14.99) with the HiFi preset. Primary alignments were retained by excluding secondary and supplementary alignments using samtools (flags -F 0x100 -F 0x800)^5^.

For each read, alignment quality metrics were extracted from SAM tags, including the alignment score (mg) and the number of mismatches (NM). Read length was inferred from the CIGAR string, and a normalized mismatch rate was calculated as NM / read length. These metrics were compiled into a per-read summary table containing read identifier, read length, alignment score, and normalized mismatch rate, which served as the basis for haplotype assignment and downstream quality control.

Reads were assigned to haplotypes using the following criteria: 1) Uniquely aligned reads: Reads that aligned to only one haplotype assembly (maternal or paternal) were directly assigned to that haplotype. 2) Differential alignment quality: Reads that aligned to both haplotypes were assigned to the haplotype with the higher alignment score (mg), provided that the normalized mismatch rate was lower for that alignment. 3) Ambiguous reads: Reads that did not satisfy the above criteria were classified as ambiguous. Given that this class comprised only a small fraction of total reads (median phasing rate 98.75% for PacBio HiFi reads), ambiguous reads were randomly assigned to maternal or paternal haplotypes to maintain balanced coverage for downstream analyses.

After haplotype assignment, reads were realigned to their respective haplotype assemblies and used for all downstream haplotype-resolved methylation and chromatin analyses.

10 **CpG methylation and chromatin feature calling from Fiber-seq data**

PacBio Fiber-seq HiFi reads were processed to infer chromatin features and CpG methylation using a combination of PacBio kinetic signals and downstream analytical tools. Methylation of adenine residues (m6A) introduced by the Hia5 methyltransferase was first predicted from HiFi kinetic data using ft predict-m6a (v0.5.4)^6,7^. This step encodes m6A probabilities in the MM and ML BAM tags and annotates nucleosome footprints (nl, ns) and MTase-sensitive patches (al, as) within each read.

m6A-annotated reads were then aligned to the reference genome or individual haplotype assemblies using pbmm2 (v1.14.99) with the HiFi preset. CpG methylation (5mC) levels were subsequently extracted from aligned BAM files using pb-CpG-tools (v2.3.2), employing the pileup-based calling model to generate CpG methylation scores at single-base resolution.

Fiber-seq Inferred Regulatory Elements (FIREs), representing regions protected from MTase accessibility by protein binding or chromatin compaction, were identified at single-molecule resolution using ft fire^6,7^. All Fiber-seq processing steps were implemented within a Snakemake workflow to ensure reproducibility and consistency across samples.

11 **Coverage uniformity and centromeric representation**

To assess coverage uniformity and ensure adequate representation of centromeric regions, haplotype-phased reads were realigned to their corresponding maternal or paternal assemblies. Genome-wide sequencing depth was quantified using mosdepth^8^ (v3.1) by computing read depth in non-overlapping 10-kb bins across each assembly. Coverage distributions were examined for all chromosomes to evaluate uniformity and to confirm consistent read representation within centromeric satellite regions relative to chromosome arms. Bins overlapping regions flagged as low-confidence assembly segments were excluded from analysis.

12 **Centromeric satellite annotation and usage**

Centromeric satellite (cenSat) annotations for all haplotype-resolved assemblies were obtained from a companion study by our collaborators^1^, in which complete telomere-to-telomere centromeric repeat structures were systematically annotated. Detailed methods for repeat identification, higher-order repeat (HOR) classification, and curation of centromeric satellite subfamilies are described in that study.

In the present work, cenSat annotations were used as fixed genomic features to stratify epigenomic analyses. We adopted the same classification scheme defined in the companion study, including active higher-order repeat arrays (active_HOR), canonical and divergent HORs (HOR and dHOR), monomeric and mixed α-satellite arrays, pericentromeric satellites (HSat1A, HSat1B, HSat2, HSat3, β-satellite, and γ-satellite), and the centromere transition (ct) region.

All downstream analyses—including identification of centromeric dip regions (CDRs), quantification of DNA methylation and chromatin features, and assessment of mutation enrichment—were performed by intersecting haplotype-resolved epigenomic data with these predefined cenSat annotations. No modifications to cenSat boundaries or classifications were introduced in this study.

To restrict analyses to bona fide centromeric and pericentromeric regions, cenSat annotations were further filtered based on genomic context. Specifically, satellite annotations located outside annotated centromeric or pericentromeric domains—such as rare α-satellite or satellite fragments detected in subtelomeric or telomere-adjacent regions—were excluded. Only cenSat elements overlapping or immediately flanking annotated centromeric domains were retained. This filtering ensured that downstream epigenomic and mutational analyses focused exclusively on centromere-proximal satellite DNA and were not confounded by isolated satellite occurrences elsewhere in the genome.

13 **Identification of centromeric dip regions (CDRs)**

Centromeric dip regions (CDRs) were identified within active higher-order repeat (active_HOR) arrays based on local CpG hypomethylation patterns. For each active_HOR array, CpG methylation levels were first summarized in non-overlapping 5-kb bins using haplotype-resolved methylation calls. Active_HOR arrays that did not exhibit any discernible hypomethylated troughs were excluded from further CDR analysis.

For active_HOR arrays containing candidate hypomethylated regions, CpG methylation was recalculated at higher resolution using non-overlapping 1-kb bins. Within each array, local methylation profiles were examined to identify contiguous regions exhibiting substantially reduced methylation relative to the surrounding HOR background. Regions showing methylation levels within the lowest 10% of values across the active_HOR array were considered candidate CDRs.

CDR boundaries were refined by visual inspection of the high-resolution methylation profiles to ensure that each identified region corresponded to a single, well-defined hypomethylated domain with clear transitions to flanking hypermethylated HOR sequence. This step was used to exclude fragmented or ambiguous troughs and to define consistent CDR boundaries across haplotypes.

For visualization and comparative analyses, methylation profiles were plotted using 1-kb bins with a sliding-window smoothing of 5 bins. All CDR analyses were performed independently for each haplotype and cell type. Bin sizes and smoothing parameters were selected to balance spatial resolution and robustness to local noise and were applied consistently across all analyses.

14 **Quantification of CDR hypomethylation (MARS)**

To quantify the overall extent of CpG hypomethylation within each centromeric dip region (CDR), we defined a methylation area score (MARS) that integrates both the depth and breadth of hypomethylation across the region.

For each CDR, CpG methylation levels were summarized in non-overlapping 1-kb bins. Bin-level methylation values within the CDR were ranked from highest to lowest. To establish a robust reference for local hypermethylation and to minimize the influence of extreme values, the mean methylation level of the top 5% highest-methylated bins was calculated and used as an internal baseline.

To further reduce sensitivity to outliers, bins corresponding to the top and bottom 5% of methylation values were excluded from subsequent integration. The remaining middle 90% of bins were used to quantify the hypomethylated area. For each retained bin, the difference between the reference methylation level (top 5% mean) and the bin’s methylation value was calculated. These differences were then summed across all retained bins to yield the methylation area score (MARS) for that CDR.

This formulation captures the cumulative hypomethylated capacity of the CDR while reducing sensitivity to local noise or extreme bin-level fluctuations. MARS values were computed independently for each haplotype and cell type and were used for all comparative analyses of CDR hypomethylation across inheritance and cell-fate transitions. Bin sizes and smoothing parameters were selected to balance spatial resolution and robustness to local noise and were applied consistently across all analyses.

15 **Quantification of FIRE signal and FIRE area score (FAS)**

Fiber-seq Inferred Regulatory Elements (FIREs) were quantified using the output of the FIRE pipeline^6,7^. Following execution of the FIRE workflow, chromatin state annotations were obtained from the all_element_coverages.bed file, which reports the fractional coverage of FIRE, linker, and nucleosome states for each genomic interval. To quantify FIRE signal within centromeric regions, these annotations were intersected with centromeric satellite (cenSat) coordinates. FIRE, linker, and nucleosome coverage values were aggregated into non-overlapping 2-kb bins, and mean coverage values for each state were calculated within each bin.

For each bin, a normalized FIRE signal was defined as the proportion of FIRE coverage relative to the total chromatin signal: FIRE signal = FIRE / (FIRE + Linker + Nucleosome). This normalization controls for local coverage differences and enables direct comparison across regions and samples. For visualization, FIRE signal profiles were smoothed using a sliding window of five consecutive bins.

To quantify cumulative protein occupancy within centromeric dip regions (CDRs), we defined a FIRE area score (FAS) using a strategy analogous to that used for the methylation area score (MARS). Briefly, FIRE signal values within each CDR were ranked, and the bottom 5% of bins with the lowest FIRE signal were used to establish a reference level. To reduce sensitivity to outliers, the top and bottom 5% of bins were excluded, and the remaining middle 90% of bins were retained for integration. For each retained bin, the difference between the reference FIRE signal and the bin’s FIRE signal was calculated, and these differences were summed across the CDR to yield the FIRE area score (FAS). FAS values were computed independently for each haplotype and cell type and were used for quantitative comparisons of centromeric protein occupancy across inheritance and cell-fate transitions.

16 **Nucleosome footprint identification and size classification**

Nucleosome footprints were identified from haplotype-phased Fiber-seq reads aligned to the corresponding maternal or paternal assemblies. These alignments contained precomputed nucleosome annotations encoded in the ns (nucleosome start) and nl (nucleosome length) tags, which were generated during Fiber-seq processing using fibertools.

For each aligned read, nucleosome positions and footprint lengths were extracted based on the ns and nl tags, allowing single-molecule–resolved identification of nucleosome occupancy and size. Nucleosome footprint lengths were then aggregated across reads to generate genome-wide and centromere-specific nucleosome size distributions.

To quantify nucleosome organization within centromeric regions, active higher-order repeat (active_HOR) arrays were segmented into non-overlapping 10-kb bins. Within each bin, nucleosome footprints were classified by size, and the number of nucleosomes in each size category was counted. The relative proportion of each nucleosome size class was then calculated by normalizing counts to the total number of nucleosome footprints detected within the bin. These bin-level nucleosome density and size composition profiles were used to compare chromatin architecture across haplotypes and cell types, with a particular focus on centromeric dip regions (CDRs) and flanking centromeric satellite domains.

17 **De novo mutation detection and filtering**

De novo variants arising during reprogramming and differentiation were identified using haplotype-resolved PacBio HiFi sequencing data and two complementary variant-calling strategies. Phased HiFi reads were independently aligned to their corresponding maternal or paternal assemblies, and variants were detected in peripheral blood mononuclear cells (PBMCs), induced pluripotent stem cells (iPSCs), and neural progenitor cells (NPCs).

First, variants were called separately in PBMCs, iPSCs, and NPCs using DeepVariant^9^. Variants detected in PBMCs were treated as the germline background and removed from the corresponding iPSC and NPC call sets to identify candidate reprogramming-associated events. Second, somatic variant calling was performed using GATK Mutect2^10^ in matched tumor–normal mode, with PBMCs serving as the matched control for each iPSC or NPC sample.

For both pipelines, variants were required to meet stringent filtering criteria, including a minimum sequencing depth of ≥10 reads and a variant allele fraction (VAF) ≥0.5. Variants overlapping low-confidence assembly regions flagged by Flagger or NucFlagger, as well as tandem repeat regions, were excluded. Only variants detected independently by both DeepVariant and Mutect2 were retained as high-confidence de novo mutations.

18 **Mutational signature analysis**

Single-nucleotide variants (SNVs) were stratified by genomic context into centromeric satellite (cenSat) regions and chromosome arm regions based on centromere annotations (see Genome annotation). For each region, SNVs were classified into the 96 trinucleotide substitution categories (SBS96) according to the mutated base and its immediate 5′ and 3′ flanking nucleotides, following the COSMIC convention. Mutation counts were normalized to relative frequencies within each region.

Overall similarity between the mutational spectra of chromosome arms and cenSat regions was quantified using cosine similarity. To assess differences in individual substitution classes, mutation frequencies were aggregated by substitution type (e.g., T>A, T>G, T>C), and fold enrichment in cenSat relative to chromosome arms was calculated. Statistical significance was evaluated using two-sided tests based on resampling of mutations within each region, with P values reported as indicated.

To infer the mutational processes underlying the observed spectra, SBS96 profiles for chromosome arms and cenSat regions were decomposed using COSMIC single base substitution signatures (v3.5). Signature exposures were estimated using non-negative least squares (NNLS) fitting, constraining all exposures to be non-negative. Prior to fitting, SBS96 profiles were normalized to sum to one. The quality of the reconstruction was evaluated by comparing the original and reconstructed profiles using cosine similarity. Signature exposures were reported as relative contributions for each genomic region.

19 **X chromosome and X-inactivation analyses**

X-chromosome inactivation status was inferred using haplotype-resolved CpG methylation at the XIST promoter region. Consistent with established epigenetic signatures of X inactivation, haplotypes exhibiting promoter hypermethylation were classified as active X chromosomes (Xa), whereas those with promoter hypomethylation were classified as inactive X chromosomes (Xi).

To quantify chromosome-arm–level DNA methylation, chromosome X was partitioned into non-overlapping 1-kb bins, and mean CpG methylation was calculated for each bin. Bins overlapping annotated centromeric satellite regions (cenSat) were excluded. Average methylation levels across the remaining bins were computed to represent global arm-level methylation for each haplotype and cell type.

20 **Statistical analyses**

Unless otherwise specified, statistical analyses were performed using two-sided, non-parametric tests to accommodate non-normal distributions typical of epigenomic measurements. Comparisons between two groups were conducted using the Wilcoxon rank-sum test (unpaired) or Wilcoxon signed-rank test (paired), as indicated in the figure legends. For comparisons across more than two groups, pairwise Wilcoxon tests were applied.

P values < 0.05 were considered statistically significant unless otherwise stated. Exact P values are reported in the text or figure legends where relevant. For analyses involving multiple hypothesis testing, P values were adjusted using the Benjamini–Hochberg false discovery rate (FDR) correction. Adjusted P values (q values) < 0.05 were considered significant. Where unadjusted P values are shown, this is explicitly indicated.

21 **Correlation and permutation analyses**

Correlation analyses were performed using Pearson’s correlation coefficient unless otherwise specified. Correlations were used to assess relationships between continuous variables, including but not limited to: 1) methylation area score (MARS) across generations or cell types; 2) relationships between CDR restoration (ΔMARS) and active HOR demethylation (ΔHOR_Meth); 3) concordance of epigenomic features across inherited haplotypes. Where appropriate, correlations were stratified by chromosome, haplotype, or cell type, as indicated in the corresponding figures.

22 **Software versions and computational environment**

All analyses were performed in a Linux-based high-performance computing environment. Statistical analyses and data visualization were conducted primarily in R (version ≥ 4.2.0) using base R functions and commonly used packages, including ggplot2, dplyr, and stats. Genomic interval operations were performed using bedtools^11^ (v2.30.0) and custom Python and shell scripts. Specific software versions for sequencing alignment, methylation calling, and Fiber-seq analyses are detailed in the corresponding Methods sections.

23 **Ethics statement**

All human samples used in this study were derived from a previously described three-generation pedigree recruited in St. Louis, Missouri. All participants provided written informed consent for open data sharing, broad research use, and induced pluripotent stem cell (iPSC) derivation, as described in the companion study by the Miga laboratory. Sample collection and generation of lymphoblastoid cell lines and iPSC lines were conducted under protocols approved by the appropriate institutional review boards. The present study analyzed de-identified, previously generated genomic and epigenomic data and did not involve additional participant recruitment or intervention.

24 **Code availability**

Custom scripts used for CDR, MARS, and FAS analyses are available from the authors upon reasonable request.

25 **Data Availability**
