## Supplemental Figures for "Fully T2T pedigree assemblies reveal genetic stability and epigenetic plasticity of human centromeres across inheritance and cell-fate transitions"

**Supplementary Figures**

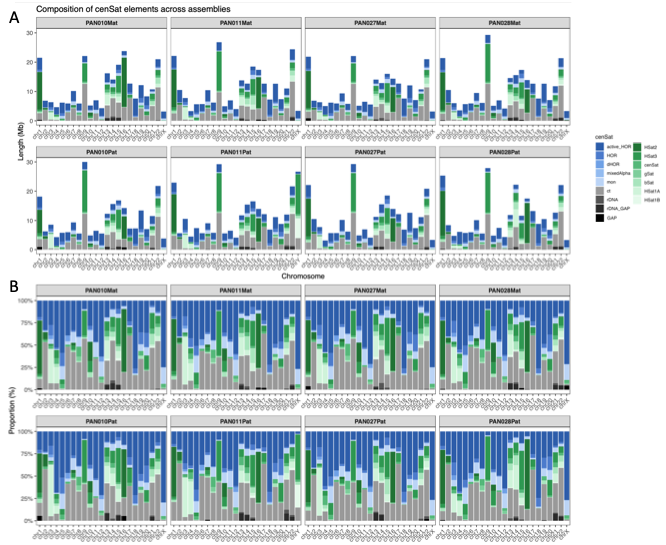

Supplementary Figure 1 | Chromosome-specific distribution of cenSat elements across assemblies. (A) Absolute length (Mb) of cenSat subcomponents on each chromosome for all maternal and paternal assemblies. (B) Relative proportion (%) of each cenSat element per chromosome.

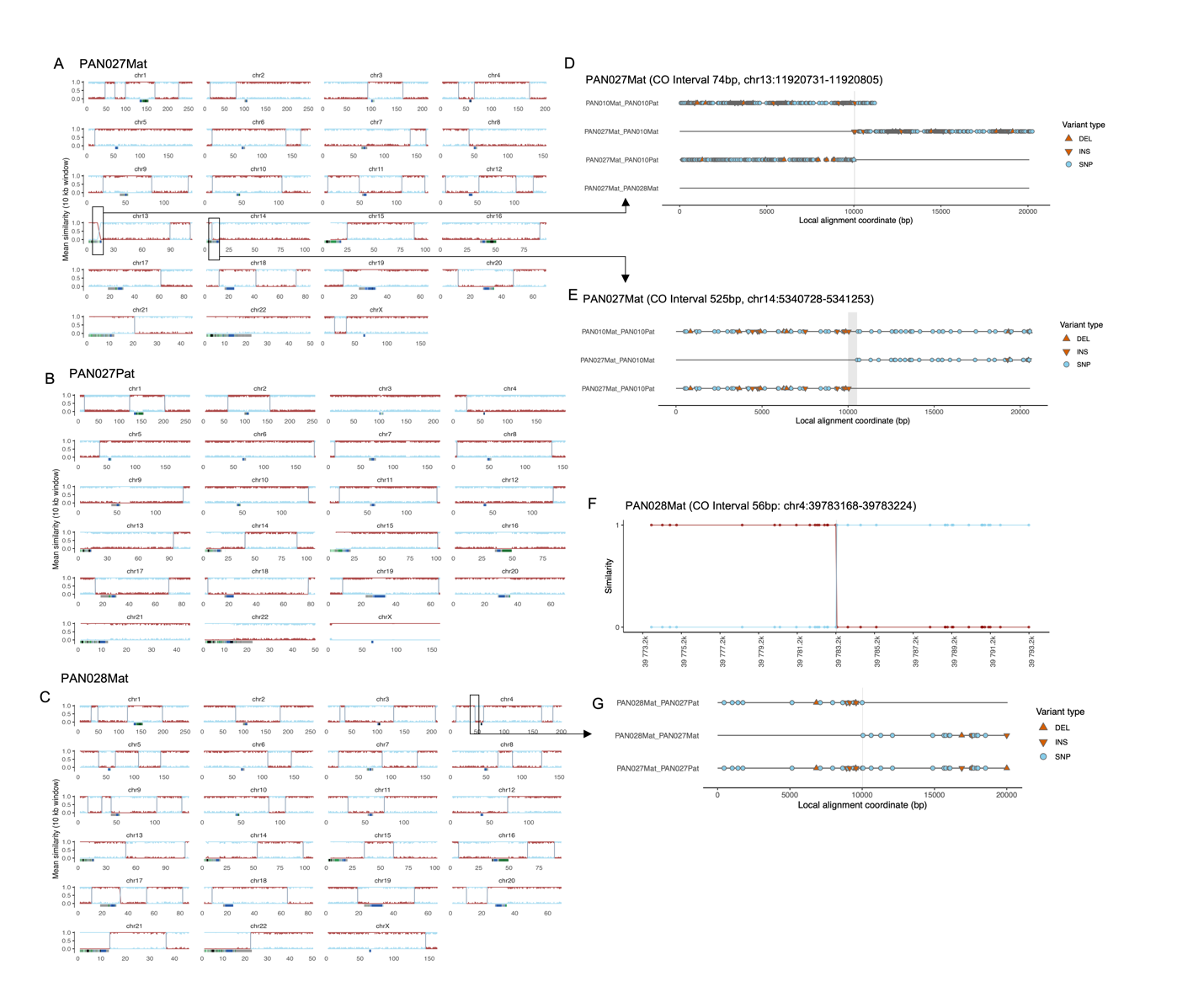

Supplementary Figure 2 | Genetic variant inferred meiotic recombination across the pedigree. For each chromosome, haplotype blocks are shown for PAN027Mat (A), PAN027Pat (B), and PAN028Mat (C). Blocks inherited from the maternal allele are shown in red, and those inherited from the paternal allele are shown in light blue. The y-axis indicates sequence similarity between each inherited block and its inferred parental haplotype, enabling the identification of meiotic crossover (CO) breakpoints. The bottom annotation track indicates centromeric satellite (cenSat) regions. D-E, Local sequence realignment validating two crossover events in acrocentric region chr13: 11,920,731–11,920,805 (interval 74bp) (D) and chr14: 53,407,28–53,412,53 (interval 525bp) (E) in PAN027Mat. Pairwise alignment of the parental haplotypes and the inherited offspring haplotype are shown. (F) Visualization of the smallest inferred recombination interval (56 bp) identified on PAN028 chromosome 4, defined by informative genetic variants distinguishing parental haplotypes (PAN027Mat, red; PAN027Pat, light blue). Each dot represents a genetic variant supporting the breakpoint. (G) Local realignment of ±10 kb flanking sequences around the recombination interval shown in (F), confirming precise breakpoint localization. Straight lines indicate matched sequences; symbols denote genetic variants, including circles for single-nucleotide polymorphisms (SNPs), triangles for deletions (DEL), and inverted triangles for insertions (INS).

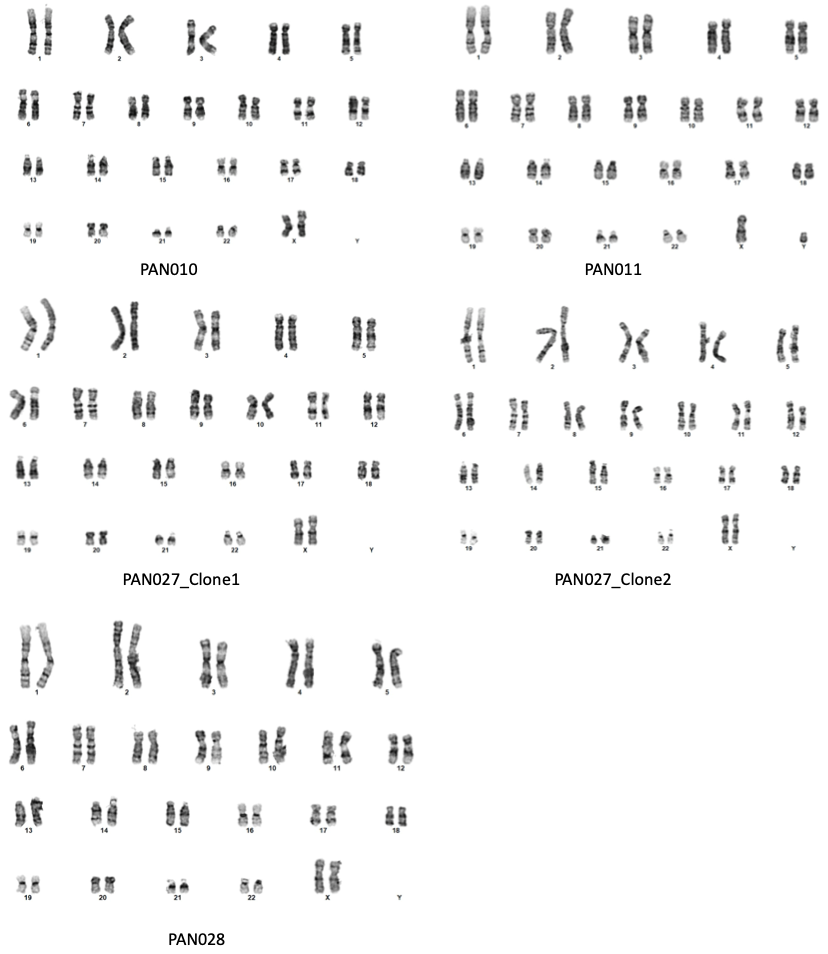

Supplementary Figure 3 | Karyotype analysis of iPSC lines. Representative GTW-banded karyotypes of each iPSC lines clones. All iPSC lines exhibited a normal karyotype at 450-band resolution. Chromosomes were counted and analyzed from 20 metaphase spreads per clone, confirming the absence of aneuploidy or large-scale structural abnormalities (>10 Mb).

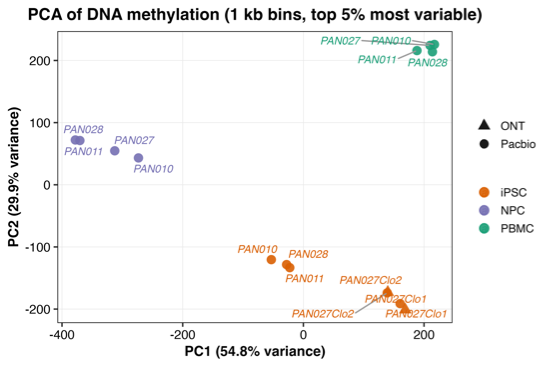

Supplementary Figure 4 | PCA of genome-wide CpG methylation across individuals, cell types, clones, and sequencing platforms. Principal component analysis (PCA) of DNA methylation (1-kb bins, top 5% most variable) for all samples.

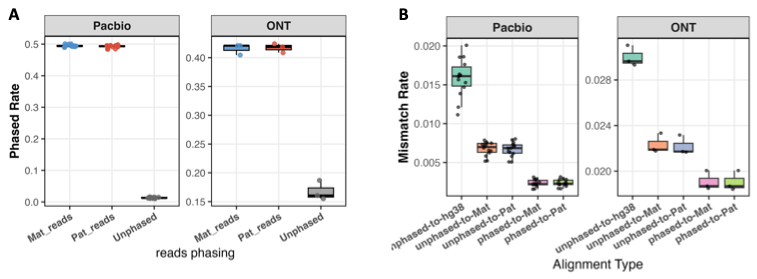

Supplementary Figure 5 | Haplotype-resolved read phasing and alignment accuracy for PacBio HiFi and ONT long reads. (A) Read-level phasing rates for PacBio HiFi (left) and ONT (right) sequencing. (B) Read mismatch rates before and after haplotype-aware phasing.

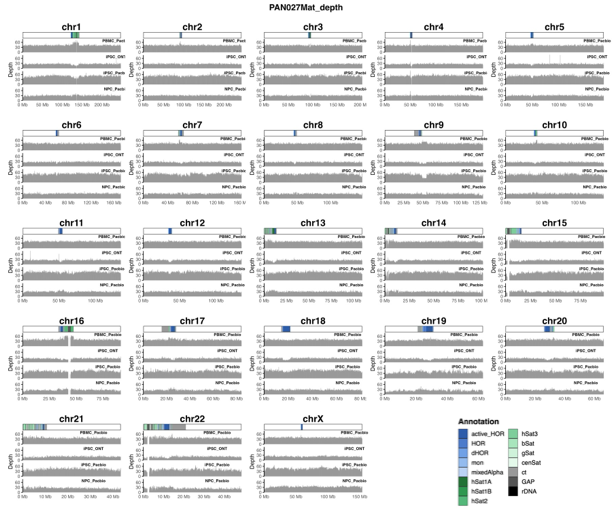

Supplementary Figure 6 | Genome-wide read-depth profile across all chromosomes in the PAN027Mat assembly. Grey bars represent read coverage calculated in 10 kb bins, with problematic (flagged) regions excluded from analysis (GAP region). Centromeric regions are indicated by highlighted bars at the top of each panel.

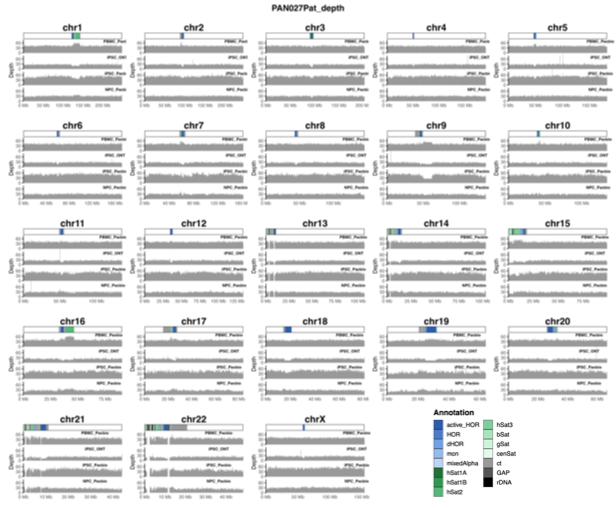

Supplementary Figure 7 | Genome-wide read-depth profile across all chromosomes in the PAN027Pat assembly. Grey bars represent read coverage calculated in 10 kb bins, with problematic (flagged) regions excluded from analysis (GAP region). Centromeric regions are indicated by highlighted bars at the top of each panel.

­­
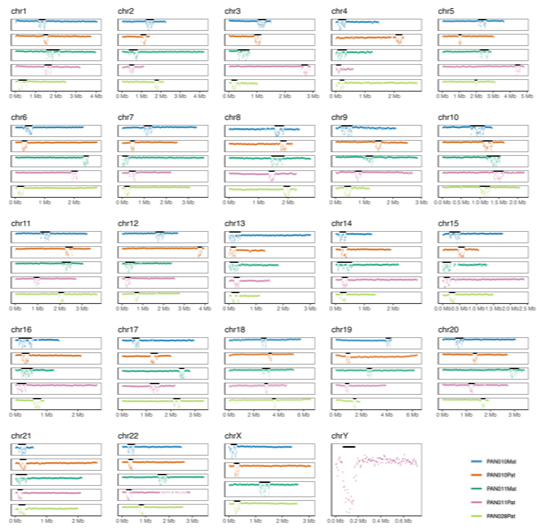

Supplementary Figure 8 | CpG methylation profiles across active HOR arrays. Methylation levels were calculated in 5-kb bins across active higher-order repeat (HOR) arrays for all chromosomes. Black bars indicate predicted centromeric dip regions (CDRs). The y-axis denotes CpG methylation level.

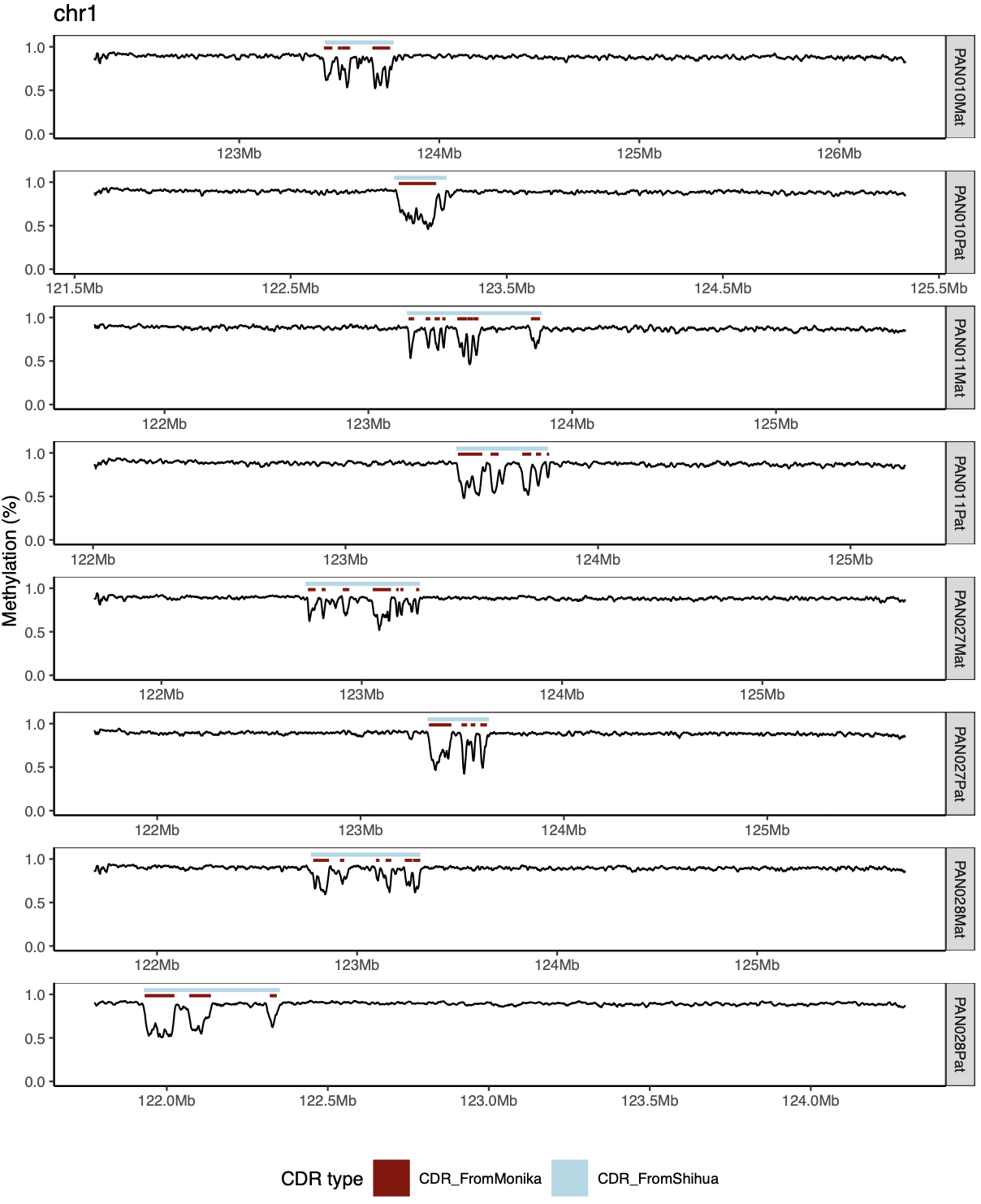

Supplementary Figure 9 | Concordance between CDR definitions derived in this study and previously reported subCDR annotations. Representative DNA methylation profiles across chromosome 1 are shown for all haplotypes. Black lines indicate 1kb binned CpG methylation levels along the active HOR array. Light blue bars denote centromeric dip regions (CDRs) identified using the approach developed in this study, whereas dark red bars indicate subCDRs identified using a previously reported method. The y-axis represents DNA methylation level.

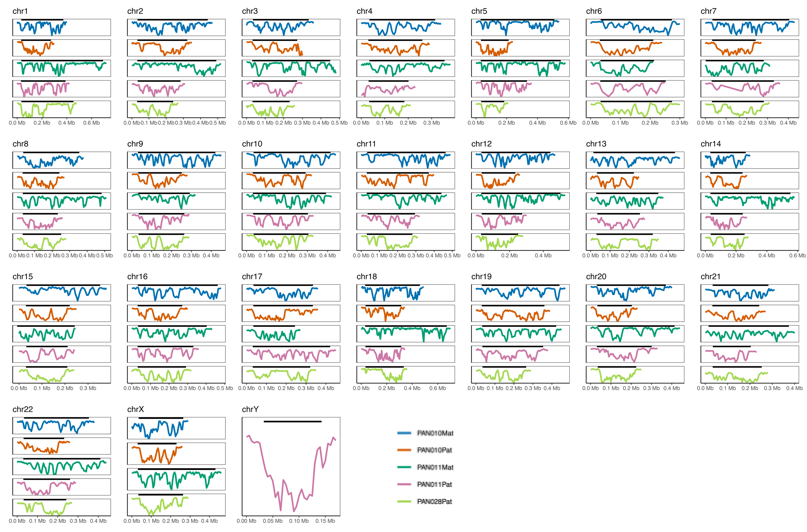

Supplementary Figure 10 | Zoomed-in CpG methylation profiles of centromeric dip regions (CDRs). Expanded views of CDR methylation patterns shown in Supplementary Fig. 8.

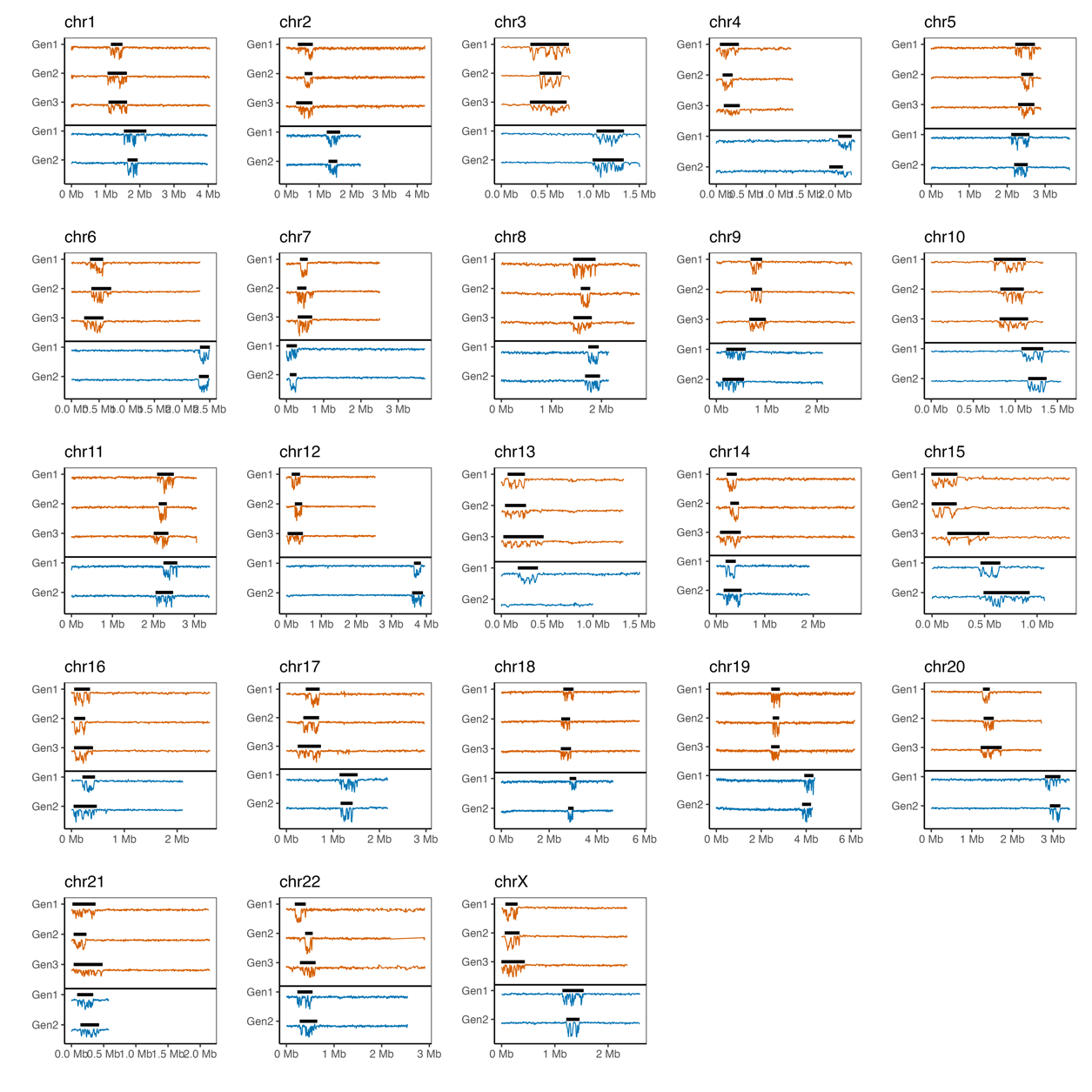

Supplementary Figure 11 | Inheritance of active HOR arrays and centromeric dip regions (CDRs) across three generations. Active higher-order repeat (HOR) arrays and their associated CDRs are shown for each chromosome across three generations (orange) and two generations (blue).

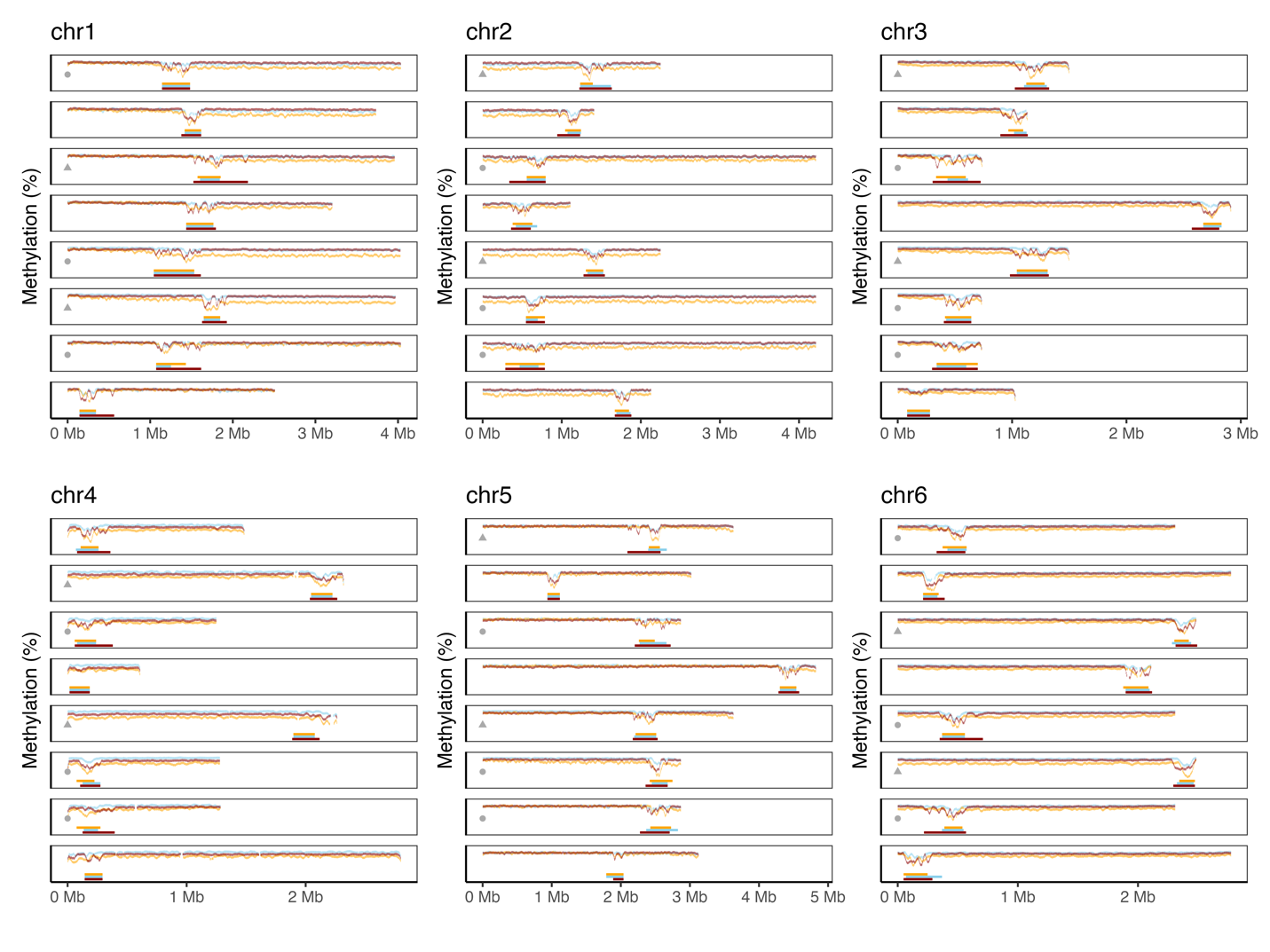

Supplementary Figure 12 | Allele-resolved active HOR methylation profiles across chromosomes 1–6. CpG methylation profiles across active HOR arrays are shown for all haplotypes and cell types. For each chromosome, panels are ordered from top to bottom as PAN010Mat, PAN010Pat, PAN011Mat, PAN011Pat, PAN027Mat, PAN027Pat, PAN028Mat, and PAN028Pat. Circles indicate active HOR arrays transmitted across three generations, whereas triangles denote two-generation transmission, and blank denotes independent haplotypes. Colors represent cell types: PBMC (brick red), iPSC (light blue), and NPC (orange).

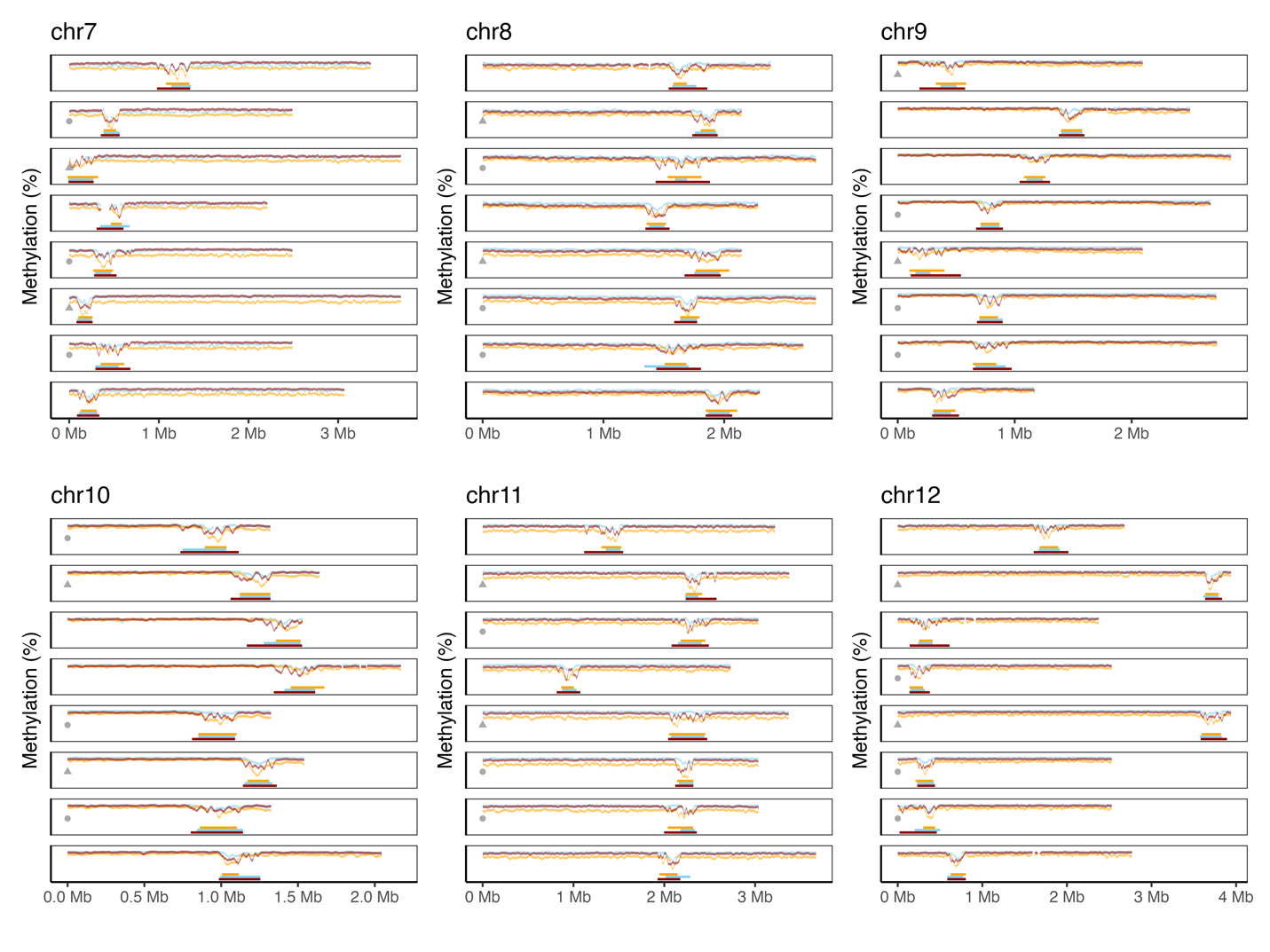

Supplementary Figure 13 | Allele-resolved active HOR methylation profiles across chromosomes 7–12. CpG methylation profiles across active HOR arrays are shown for all haplotypes and cell types. For each chromosome, panels are ordered from top to bottom as PAN010Mat, PAN010Pat, PAN011Mat, PAN011Pat, PAN027Mat, PAN027Pat, PAN028Mat, and PAN028Pat. Circles indicate active HOR arrays transmitted across three generations, whereas triangles denote two-generation transmission, and blank denotes independent haplotypes. Colors represent cell types: PBMC (brick red), iPSC (light blue), and NPC (orange).

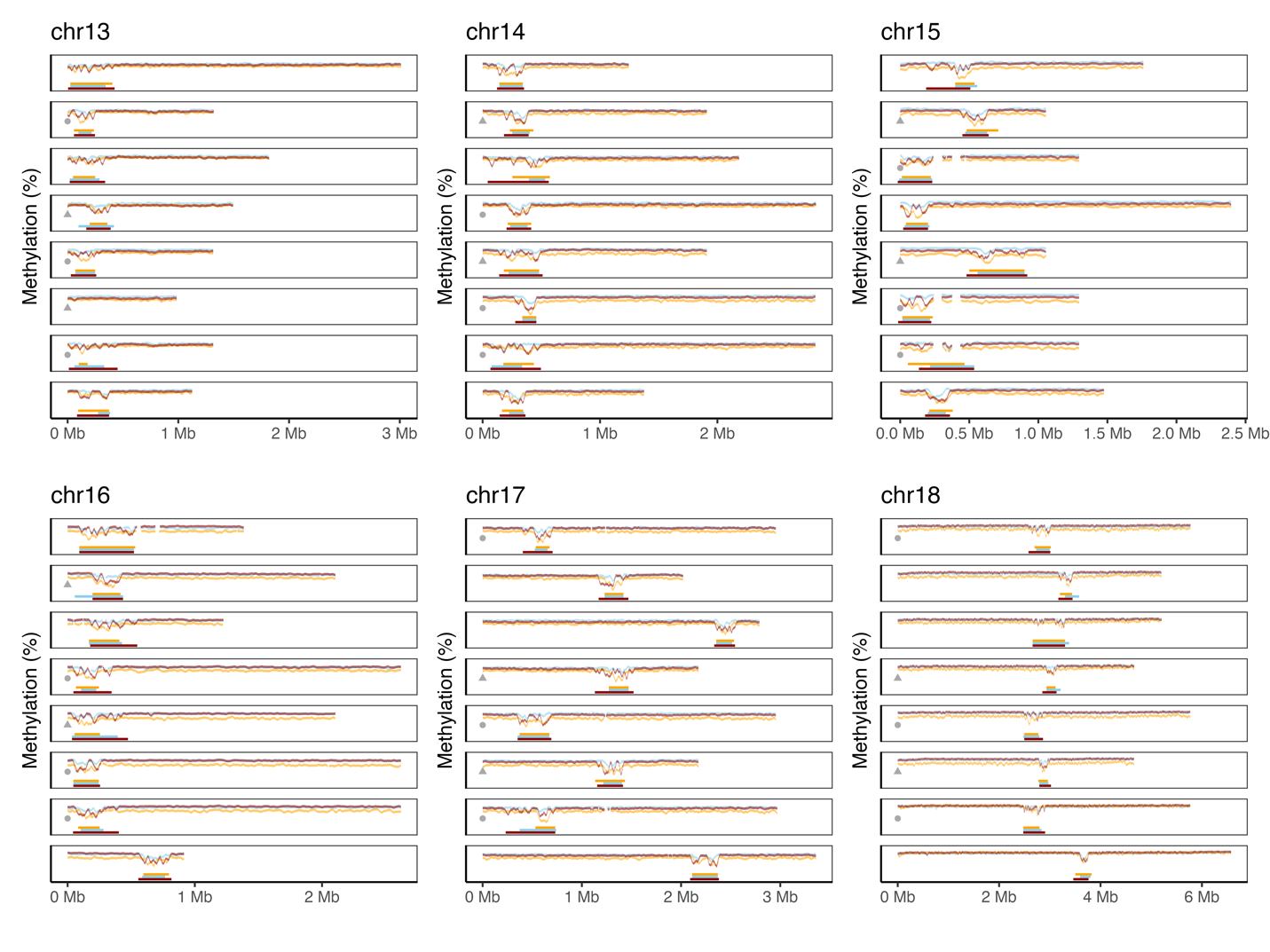

Supplementary Figure 14 | Allele-resolved active HOR methylation profiles across chromosomes 13–18. CpG methylation profiles across active HOR arrays are shown for all haplotypes and cell types. For each chromosome, panels are ordered from top to bottom as PAN010Mat, PAN010Pat, PAN011Mat, PAN011Pat, PAN027Mat, PAN027Pat, PAN028Mat, and PAN028Pat. Circles indicate active HOR arrays transmitted across three generations, whereas triangles denote two-generation transmission, and blank denotes independent haplotypes. Colors represent cell types: PBMC (brick red), iPSC (light blue), and NPC (orange).

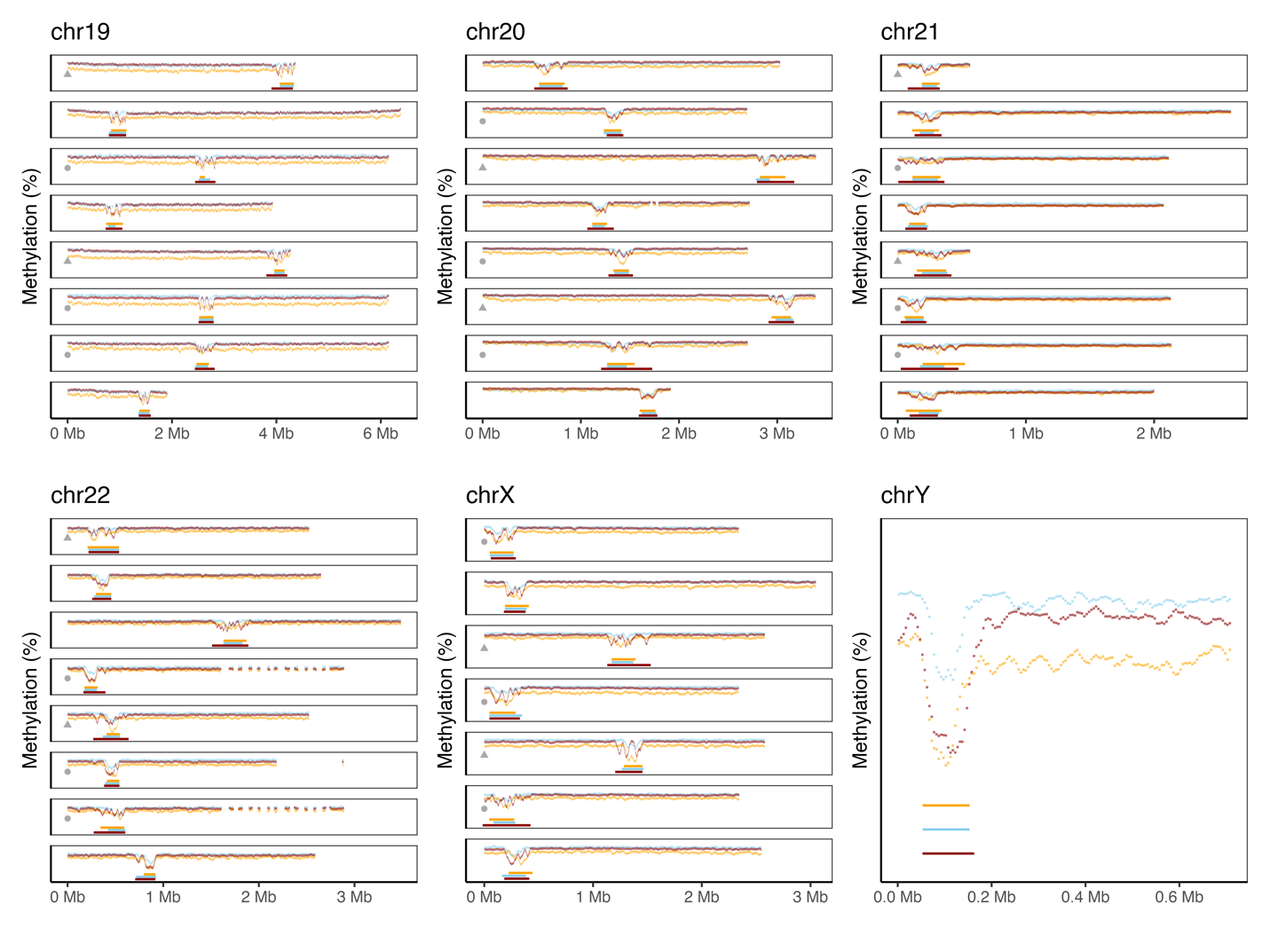

Supplementary Figure 15 | Allele-resolved active HOR methylation profiles across chromosomes 19–22, X, and Y. CpG methylation profiles across active HOR arrays are shown for all haplotypes and cell types. For each chromosome, panels are ordered from top to bottom as PAN010Mat, PAN010Pat, PAN011Mat, PAN011Pat, PAN027Mat, PAN027Pat, PAN028Mat, and PAN028Pat. Circles indicate active HOR arrays transmitted across three generations, whereas triangles denote two-generation transmission, blank denote independent haplotypes. Colors represent cell types: PBMC (brick red), iPSC (light blue), and NPC (orange).

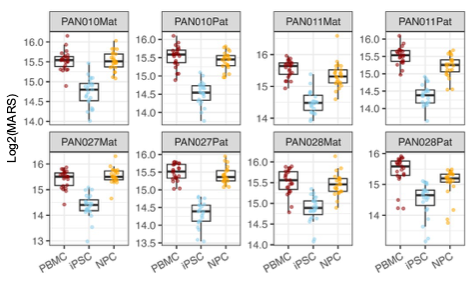

Supplementary Figure 16 | Cell-type–specific variation of CDR methylation area score (MARS) across haplotypes. Boxplots show log₂-transformed MARS values for centromeric dip regions (CDRs) across PBMCs, iPSCs, and NPCs for each haplotype.

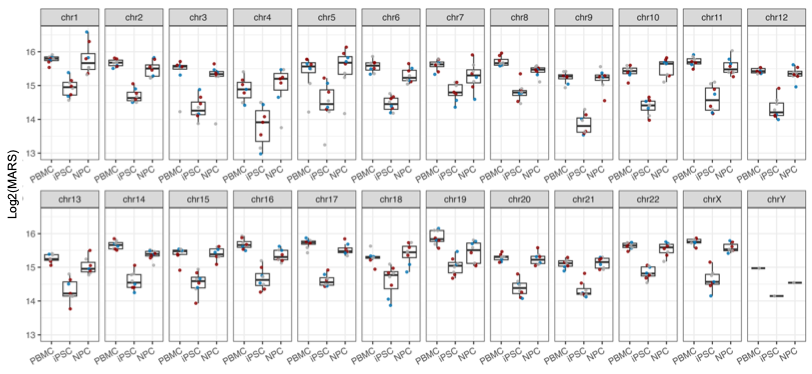

Supplementary Figure 17 | Chromosome-specific variation of CDR methylation area score (MARS) across cell types. Boxplots show log₂-transformed MARS values for centromeric dip regions (CDRs) across PBMCs, iPSCs, and NPCs, displayed separately for each chromosome.

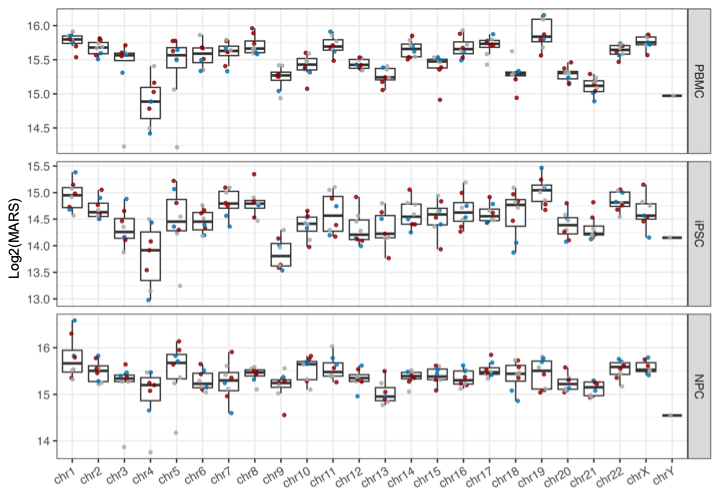

Supplementary Figure 18 | Cell-type–stratified distribution of CDR methylation area score (MARS) across chromosomes. Log₂-transformed MARS values for centromeric dip regions (CDRs) are shown for each chromosome, stratified by cell type (PBMCs, iPSCs, and NPCs).

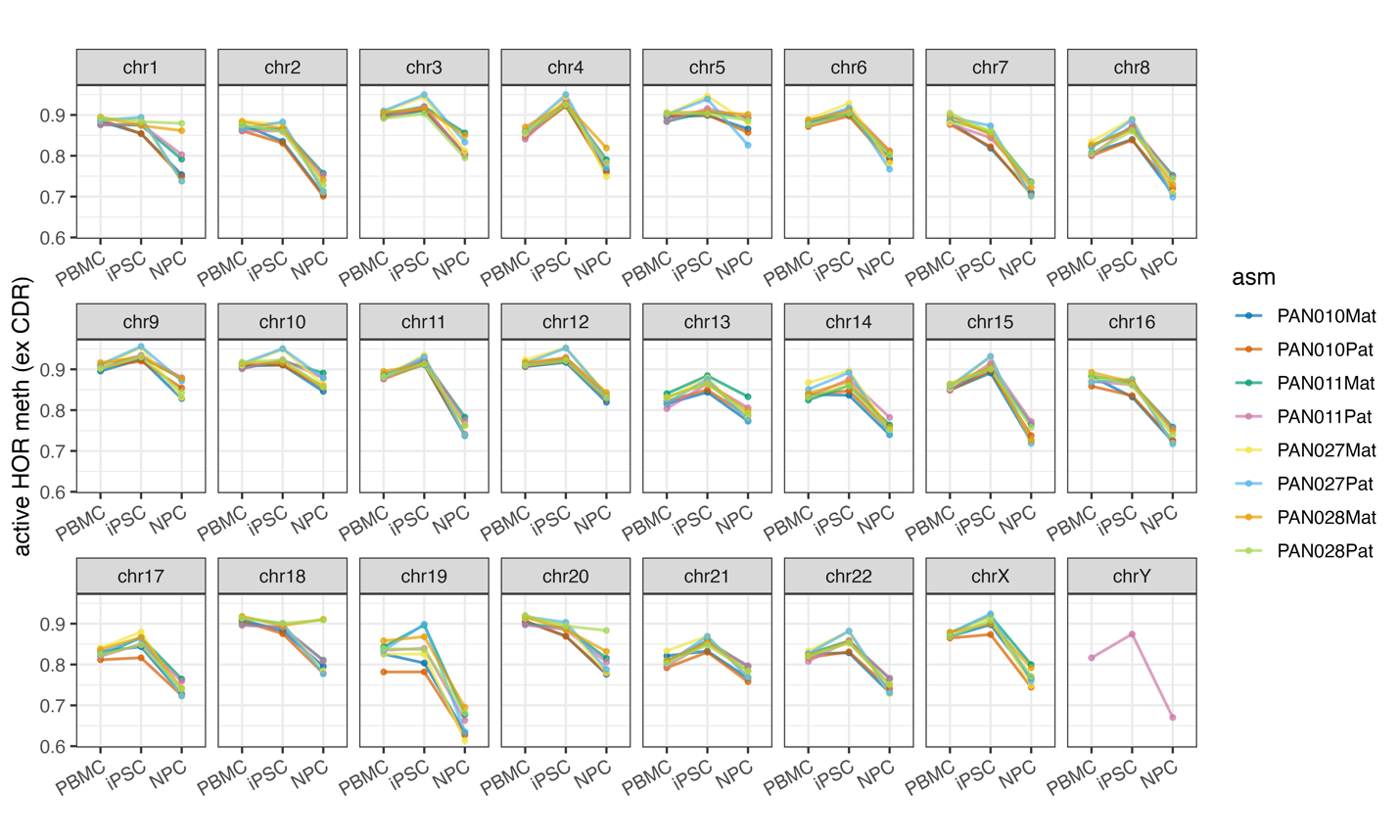

Supplementary Figure 19 | Chromosome-specific methylation dynamics of active HOR arrays across cell types. Average CpG methylation levels of active higher-order repeat (HOR) arrays are shown for each chromosome across PBMCs, iPSCs, and NPCs.

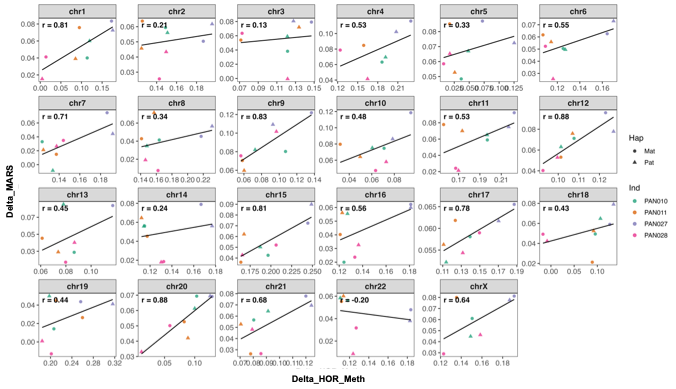

Supplementary Figure 20 | Chromosome-wise correlation between active HOR demethylation and CDR MARS recovery. Scatter plots show, for each chromosome, the correlation between changes in active HOR methylation (ΔHOR_meth) and changes in CDR methylation area score (ΔMARS) across haplotypes.

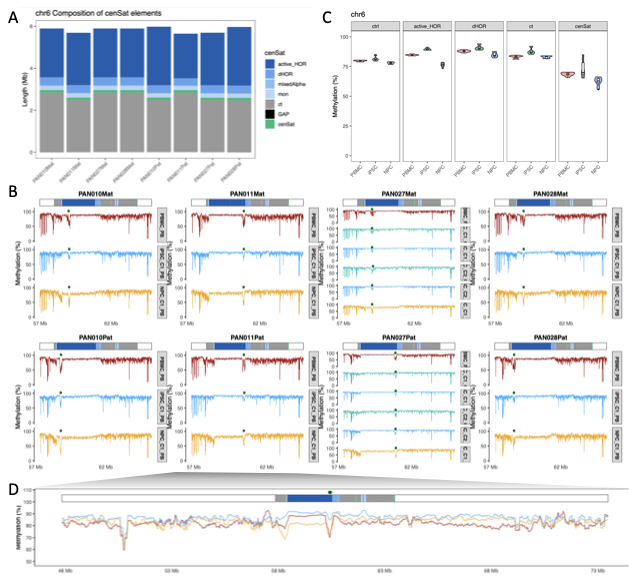

Supplementary Figure 21 | Allele-resolved structure and DNA methylation dynamics of the chromosome 6 centromere across PBMC, iPSC, and NPC lineages. A. Composition of centromeric satellites on chromosome 6. Stacked bar plot showing the length contribution of major satellite families for each haplotype assembly in the pedigree. B. Haplotype-resolved DNA methylation landscapes across the chromosome 6 centromere. For each assemblies, methylation profiles are shown for PBMC (dark red), iPSC (light blue), and NPC (gold). The top track shows the cenSat annotation, and dark-green dots denote the positions of centromeric dip regions (CDRs). C. Cell-type–specific methylation differences across major centromeric repeat classes. Violin plots (with embedded boxplots) summarizing CpG methylation for each repeat family across PBMC, iPSC, and NPC. D. Expanded view of PAN011Pat showing methylation patterns across the entire centromeric array and ±10 Mb flanking regions.

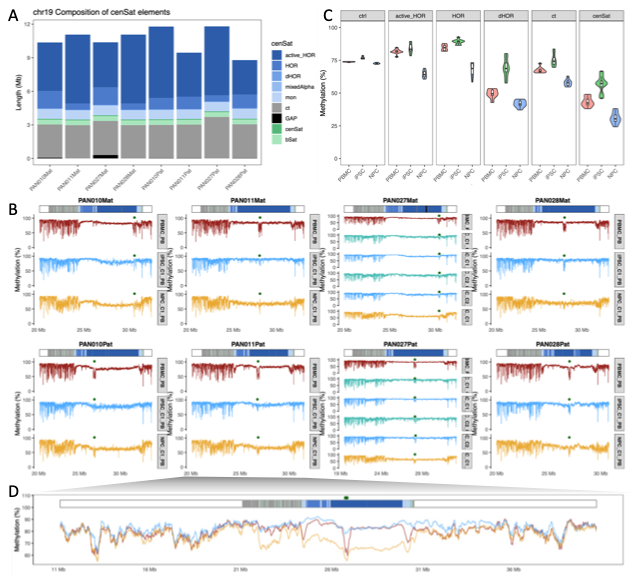

Supplementary Figure 22 | Allele-resolved structure and DNA methylation dynamics of the chromosome 19 centromere across PBMC, iPSC, and NPC lineages. A. Composition of centromeric satellites on chromosome 19. Stacked bar plot showing the length contribution of major satellite families for each haplotype assembly in the pedigree. B. Haplotype-resolved DNA methylation landscapes across the chromosome 19 centromere. For each assemblies, methylation profiles are shown for PBMC (dark red), iPSC (light blue), and NPC (gold). The top track shows the cenSat annotation, and dark-green dots denote the positions of centromeric dip regions (CDRs). C. Cell-type–specific methylation differences across major centromeric repeat classes. Violin plots (with embedded boxplots) summarizing CpG methylation for each repeat family across PBMC, iPSC, and NPC. D. Expanded view of PAN011Pat showing methylation patterns across the entire centromeric array and ±10 Mb flanking regions.

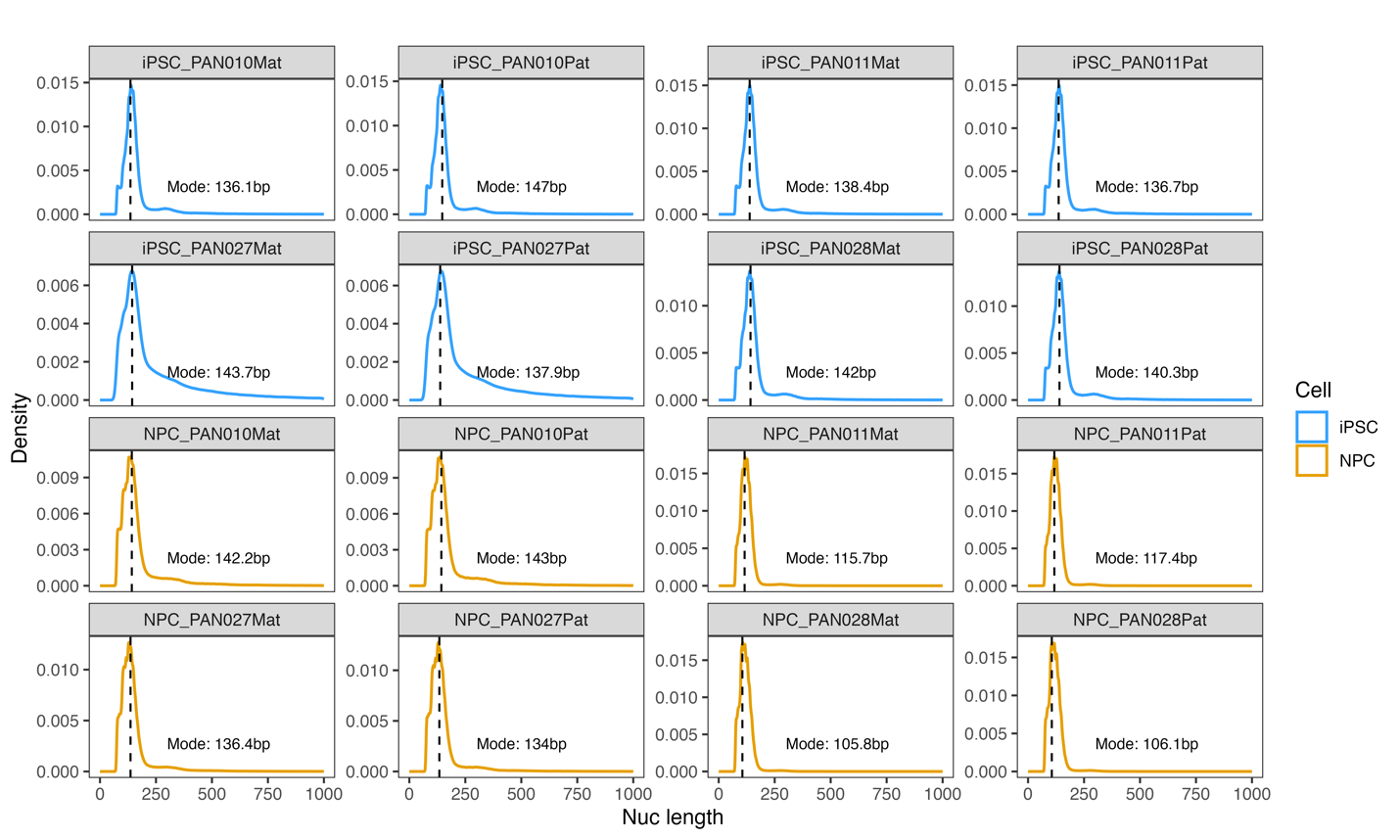

Supplementary Figure 23 | Genome-wide nucleosome footprint length distributions measured by Fiber-seq. Density plots show nucleosome footprint length distributions inferred from Fiber-seq in iPSCs (blue) and NPCs (orange) across all haplotypes. Modal nucleosome lengths are indicated for each sample.

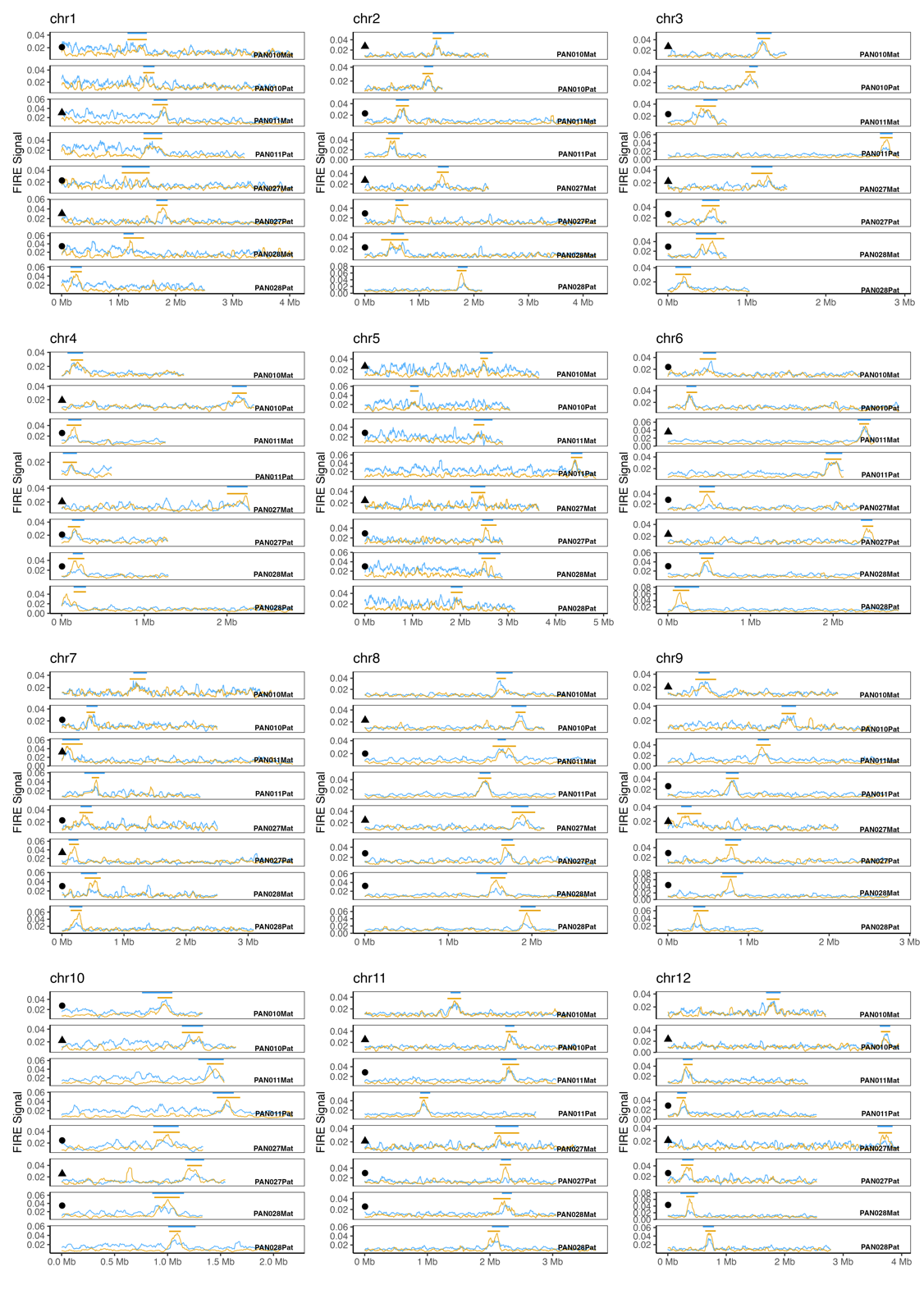

Supplementary Figure 24 | Allele-resolved FIRE signal profiles across active HOR arrays on chromosomes 1–12. Fiber-seq–derived FIRE (footprint of interacting regulatory elements) signals are shown across active higher-order repeat (HOR) arrays for all autosomes. For each chromosome, haplotypes are ordered as PAN010Mat, PAN010Pat, PAN011Mat, PAN011Pat, PAN027Mat, PAN027Pat, PAN028Mat, and PAN028Pat. Circles indicate active HOR arrays transmitted across three generations, whereas triangles denote two-generation transmission. Colors represent cell types: PBMCs (brick red), iPSCs (light blue), and NPCs (orange).

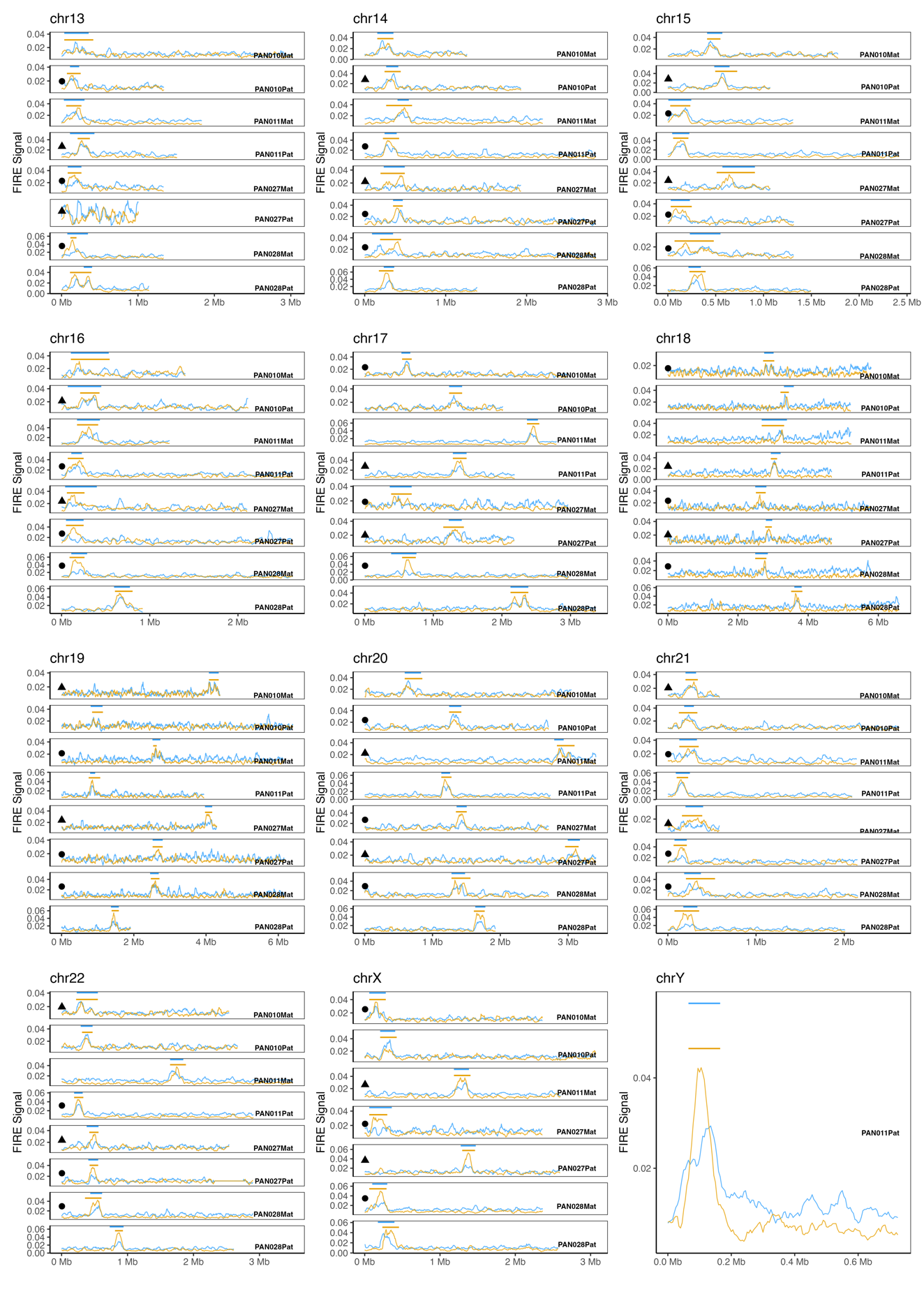

Supplementary Figure 25 | Allele-resolved FIRE signal profiles across active HOR arrays on chromosomes 13–22, X, and Y. Fiber-seq–derived FIRE (footprint of interacting regulatory elements) signals are shown across active higher-order repeat (HOR) arrays for all autosomes. For each chromosome, haplotypes are ordered as PAN010Mat, PAN010Pat, PAN011Mat, PAN011Pat, PAN027Mat, PAN027Pat, PAN028Mat, and PAN028Pat. Circles indicate active HOR arrays transmitted across three generations, whereas triangles denote two-generation transmission. Colors represent cell types: PBMCs (brick red), iPSCs (light blue), and NPCs (orange).

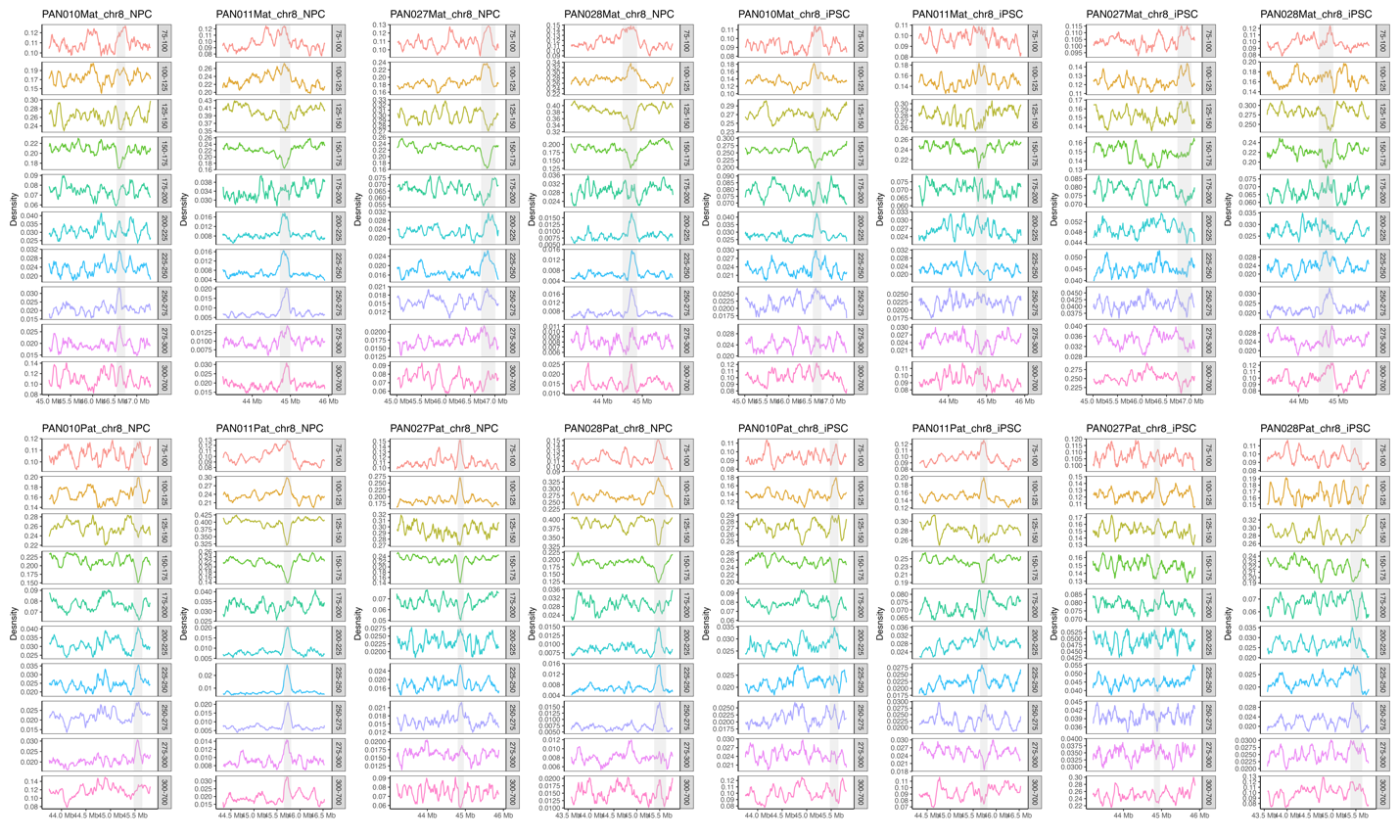

Supplementary Figure 26 | Size-resolved nucleosome footprint distributions across the chr8 active HOR array. Fiber-seq–derived nucleosome footprint density profiles are shown for the chr8 active HOR array across all haplotypes in iPSCs and NPCs. Traces are stratified by nucleosome footprint size classes, as indicated on the right.

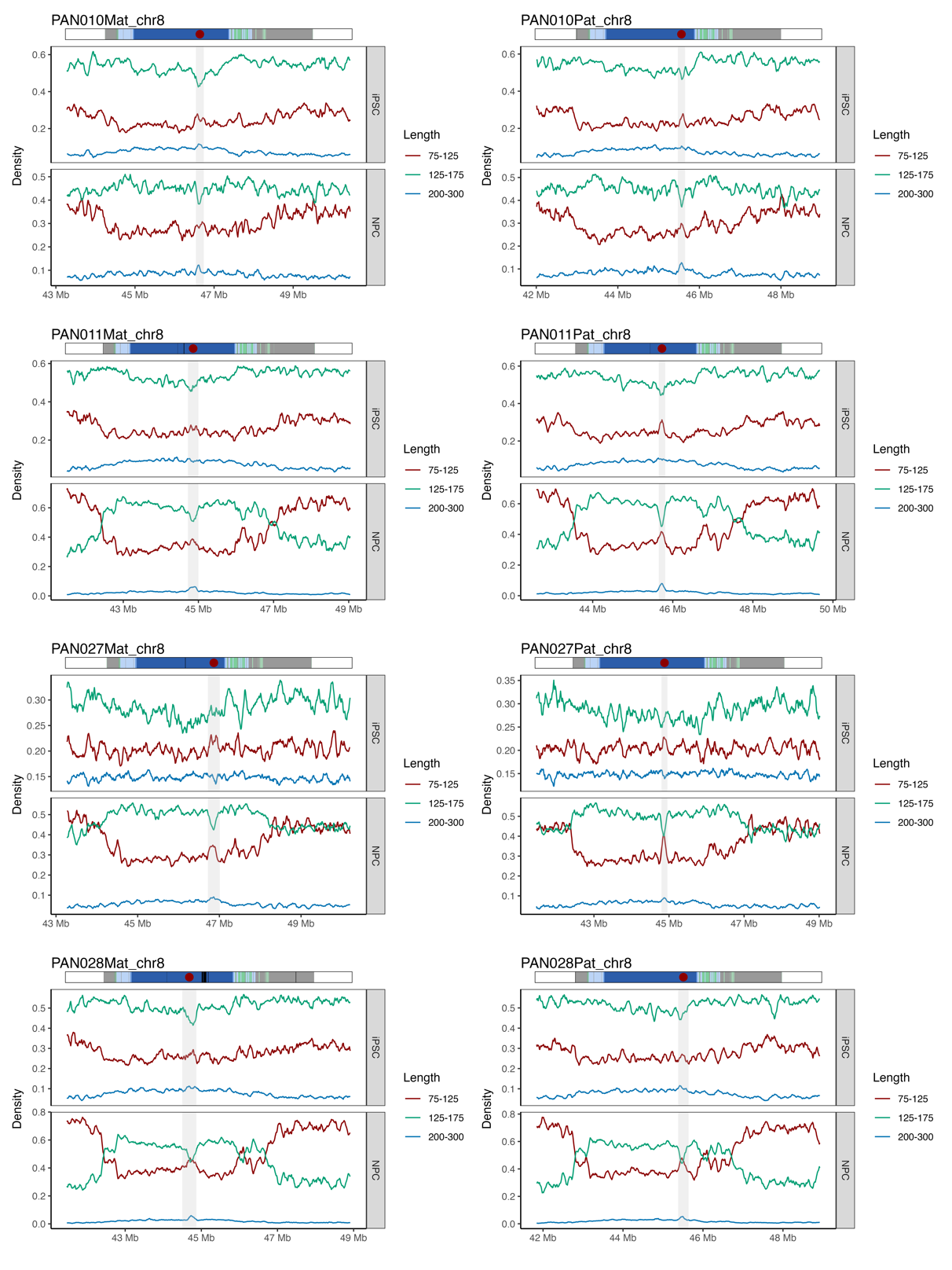

Supplementary Figure 27 | Size-resolved nucleosome footprint densities across the chr8 centromeric satellite (cenSat) domain and flanking regions. Fiber-seq–derived nucleosome footprint density profiles stratified by size classes are shown across the chr8 cenSat domain and adjacent flanking regions for all haplotypes in iPSCs and NPCs. Shaded regions indicate the cenSat interval, highlighting centromere-restricted nucleosome reorganization relative to flanking chromosomal regions.

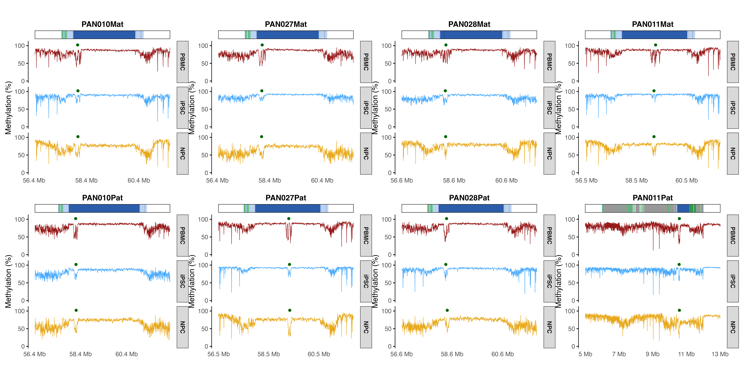

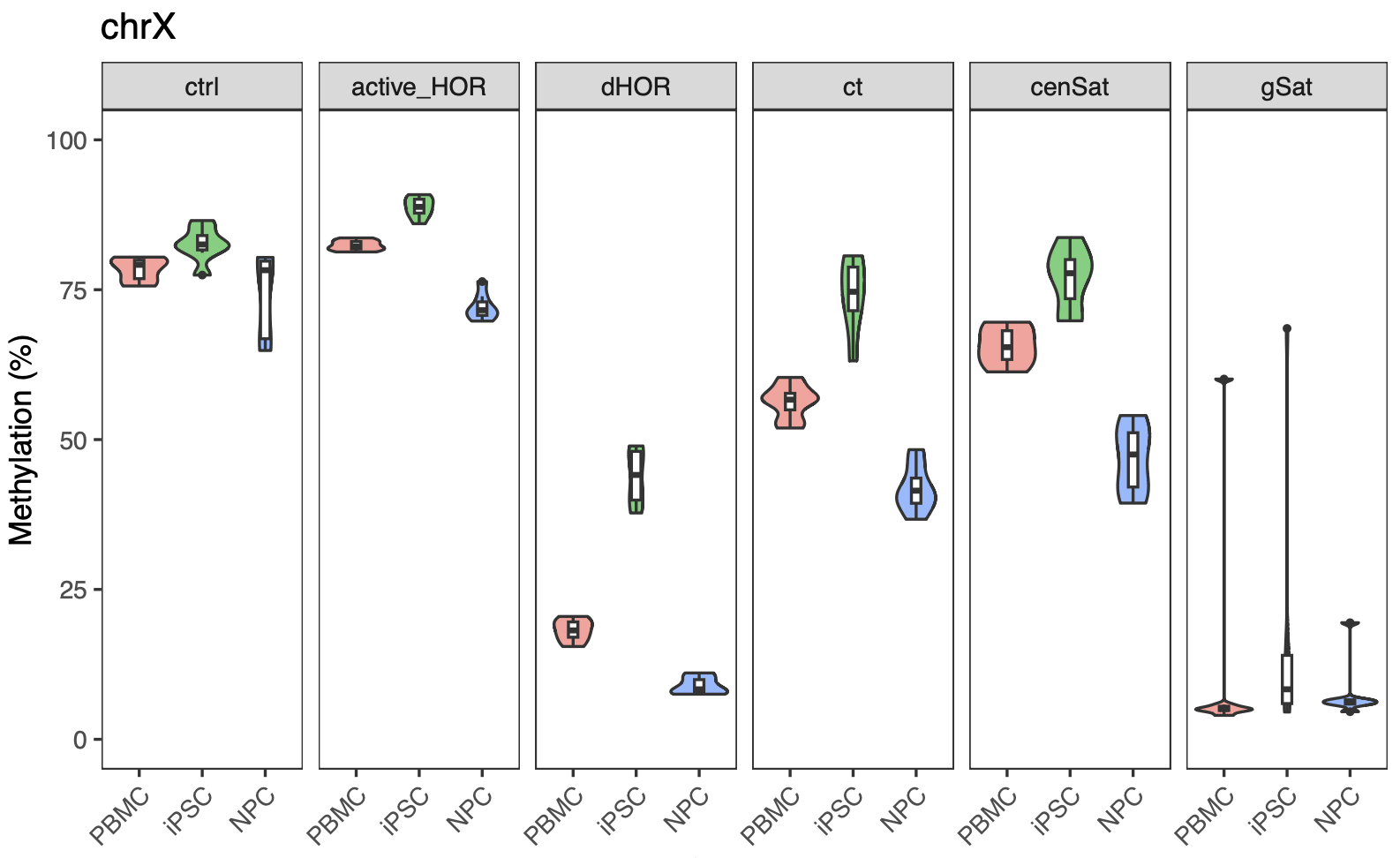

Supplementary Figure 28 | Allele-resolved structure and DNA methylation dynamics of the chromosome X centromere across PBMC, iPSC, and NPC lineages.

A. Haplotype-resolved DNA methylation landscapes across the chromosome X centromere. For each assembly, methylation profiles are shown for PBMC (dark red), iPSC (light blue), and NPC (gold). Methylation was computed in non-overlapping 5-kb windows and plotted against the coordinate system of each respective assembly. The top track shows the cenSat annotation, and dark-green dots denote the positions of centromeric dip regions (CDRs). B. Cell-type–specific methylation differences across major centromeric repeat classes. Violin plots (with embedded boxplots) summarizing CpG methylation for each repeat family across PBMC, iPSC, and NPC.

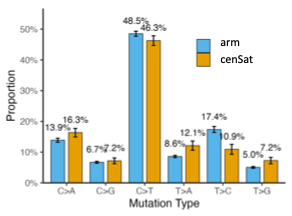

Supplementary Figure 29 | Distribution of base substitution classes in chromosome arms and centromeric satellite regions. Proportions of single-nucleotide substitution types (C>A, C>G, C>T, T>A, T>C, and T>G) in chromosome arm regions (blue) and centromeric satellite regions (cenSat; orange). Percentages above bars indicate the relative contribution of each substitution class within the corresponding genomic region.

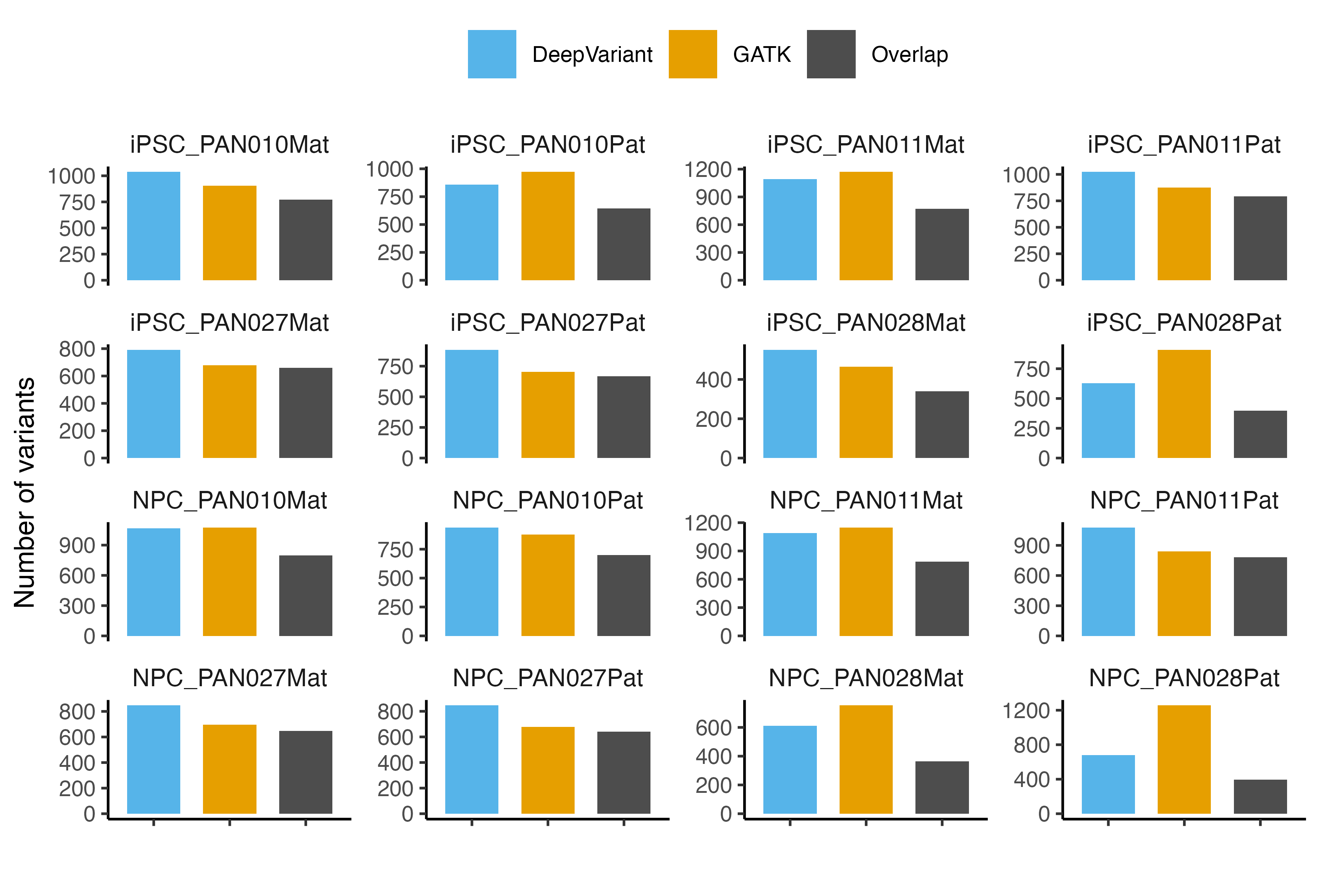

Supplementary Figure 30. Concordance of high-allele-frequency de novo variants between variant callers

Comparison of high-allele-frequency (VAF ≥ 0.5) de novo variants identified by DeepVariant and GATK across iPSC and NPC samples. Bar plots show, for each haplotype-resolved sample, the total number of variants detected by DeepVariant, GATK, and the subset supported by both callers (overlap). Variants were filtered using identical genomic masks and quality thresholds prior to comparison.

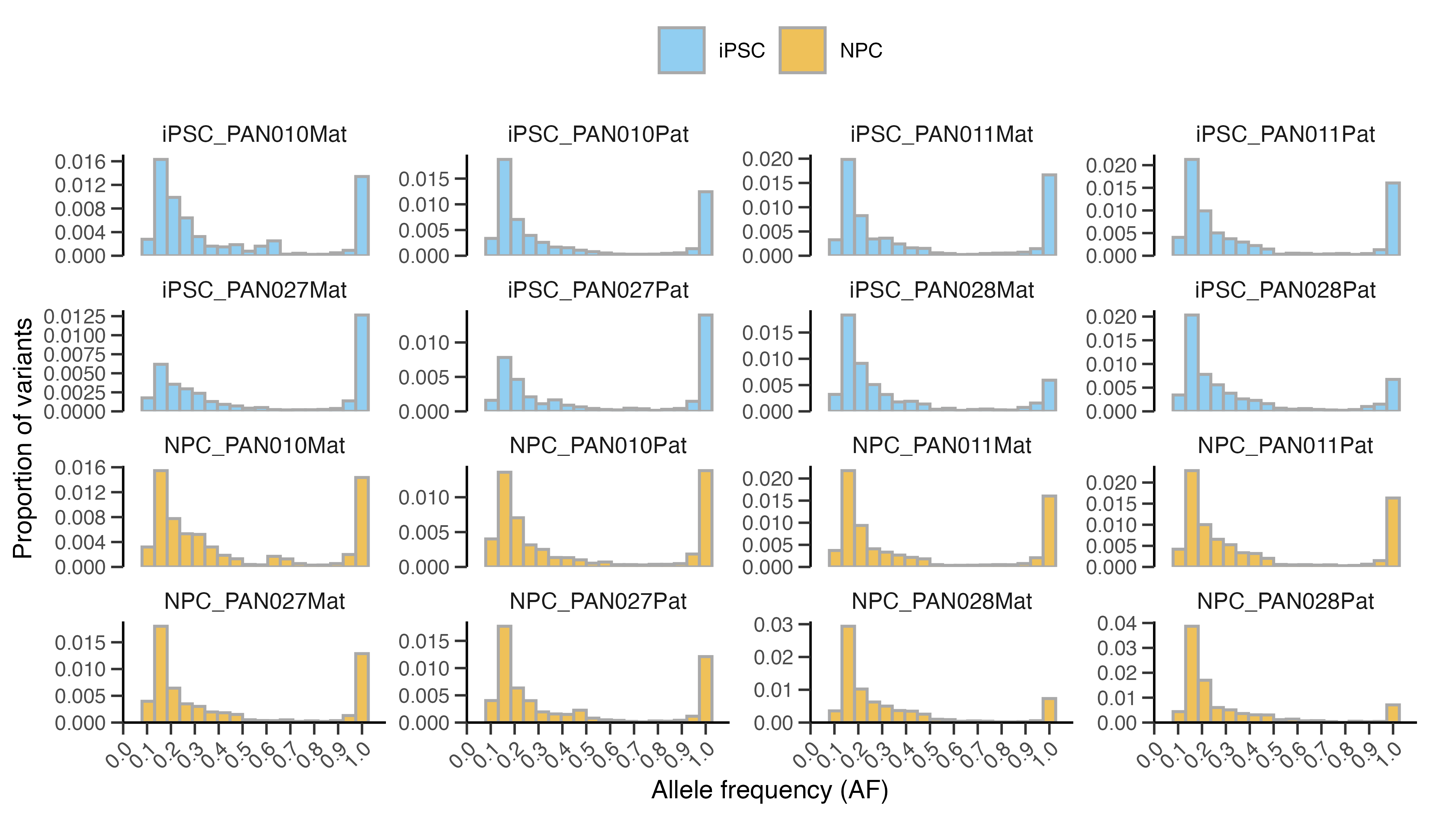

Supplementary Figure 31. Allele frequency distributions of de novo variants across samples. Histograms showing the allele frequency (VAF) distributions of de novo variants identified by DeepVariant in iPSC and NPC samples. For each haplotype-resolved sample, the x-axis represents allele frequency (AF), and the y-axis indicates the proportion of variants within each bin. Distributions are shown separately for each sample to facilitate comparison of VAF patterns across individuals and cell states.

**Supplementary Note 1. Atypical de novo mutational patterns in PAN028**

During the analysis of reprogramming-associated de novo mutations, we observed that samples from individual PAN028 exhibited distinct mutational features compared with other members of the pedigree.

First, high-allele-frequency variants (VAF ≥ 0.5), which are expected to represent early-arising or clonally expanded mutations, showed reduced concordance between DeepVariant and GATK in PAN028 relative to other samples (Supplementary Fig. 30). Across non-PAN028 samples, the overlap between the two callers accounted for 70.6–85.7% of DeepVariant calls and 66.0–97.8% of GATK calls. In contrast, PAN028 showed lower but still substantial concordance, with 58.2–63.4% of DeepVariant calls and 31.4–73.1% of GATK calls overlapping between callers. Importantly, despite this reduction, the majority of high-frequency variants in PAN028 were supported by both pipelines, indicating that variant detection remained reproducible across callers.

Second, the allele frequency distributions of de novo variants in PAN028 were markedly shifted toward lower VAF values, with a pronounced depletion of high-frequency variants compared with other individuals (Supplementary Fig. 31). In non-PAN028 samples, high-VAF variants (VAF ≥ 0.5) comprised on average 34.9% of detected de novo variants (range: 28.3–47.7%), whereas in PAN028 this proportion was substantially reduced (mean: 18.6%, range: 15.0–21.7%). This shift toward lower allele frequencies was consistently observed in both iPSC and NPC derivatives.

Together, these observations indicate that PAN028 harbors an atypical mutational landscape characterized by reduced representation of clonally stable variants and increased mutational heterogeneity. As the primary aim of this study is to characterize mutational processes associated with cellular reprogramming and their inheritance during early differentiation, analyses in the main text focus on samples exhibiting robust and concordant high-frequency mutational signals. PAN028 was therefore excluded from aggregate analyses of reprogramming-associated mutagenesis but is reported here for completeness. The underlying biological basis for this pattern remains unclear and may reflect individual-specific factors, such as increased somatic mosaicism or altered genome stability in PAN028. These possibilities were not investigated further in the present study.
